# Insights of the role of estrogen in obesity from two models of ERα deletion

**DOI:** 10.1101/2022.03.01.482291

**Authors:** Rocío del M. Saavedra-Peña, Natalia Taylor, Matthew S. Rodeheffer

## Abstract

Sex hormones play a pivotal role in physiology and disease. Estrogen, the female sex hormone, has been long implicated in having protective roles against obesity. However, the direct impact of estrogens in white adipose tissue (WAT) function and growth are not understood. Here, we show that deletion of estrogen receptor alpha (ERα) from adipocytes using *Adiponectin-cre* does not affect adipose mass in male or female mice under normal or high-fat diet (HFD) conditions. However, loss of ERα in adipocyte precursor cells (APs) via *PdgfRα-cre* leads to exacerbated obesity upon HFD feeding in both male and female mice, with subcutaneous adipose (SWAT)-specific expansion in male mice. Further characterization of these mice revealed infertility and increased plasma levels of sex hormones, including estradiol in female mice and androgens in male mice. These findings compromise the study of estrogen signaling within the adipocyte lineage using the *PdgfRα-cre* strain. However, AP transplant studies demonstrate that the increased AP hyperplasia in male SWAT upon *PdgfRα-cre*-mediated ablation of ERα is not driven by AP-intrinsic mechanisms, but are rather mediated by off-target effects. These data highlight the inherent difficulties in studying models that disrupt the intricate balance of sex hormones. Thus, better approaches are needed to study the cellular and molecular mechanisms of sex hormones in obesity and disease.

## Introduction

Obesity is defined as the excessive accumulation of WAT. The role of sex hormones in obesity and disease has been recognized for decades (Cooke and Naaz 2004; Jones, et al. 2000; Marin and Arver 1998; Seidell, et al. 1990). Sex hormones influence many aspects that can lead to obesity including food intake, energy expenditure, and the expansion of WAT (Dubuc 1985; Heine, et al. 2000; Krause, et al. 2021; Musatov, et al. 2007; Ramirez 1980; Wurtman and Baum 1980). In particular, observations in mice and humans have indicated that estrogen has an overall protective effect against obesity and metabolic disease (Andersson, et al. 1997; Stubbins, et al. 2012). After menopause, when estrogen levels in plasma decline, women are more prone to obesity and metabolic dysfunction (Aloia, et al. 1996; Andersson et al. 1997; Stubbins et al. 2012; Toth, et al. 2000). Estrogen replacement therapy is able to ameliorate these effects in humans (Andersson et al. 1997) and in ovariectomized (OVX) female mice (Stubbins et al. 2012). Furthermore, loss of function mutations in the aromatase gene, which is required for estrogen production (Meyer 1955; Meyer, et al. 1955), causes increased fat mass, hyperinsulinemia, elevated cholesterol, and fatty livers (Bilezikian, et al. 1998; Carani, et al. 1997; Conte, et al. 1994; Jones et al. 2000; Jones, et al. 2001; Morishima, et al. 1995). Similar effects are also seen upon treatment with aromatase inhibitors (Gibb, et al. 2016; Kauffman, et al. 2015). In addition, modulating the function of estrogen receptors impacts obesity and metabolism. Whole-body deletion of estrogen receptor alpha (ERα) but not ERβ leads to obesity in mice (Heine et al. 2000; Ohlsson, et al. 2000) while deletion of GPR30 receptor protects female mice from obesity (Wang, et al. 2016). Thus, both ERα and GPR30 influence obesity but the direct role of estrogen signaling on adipose biology remains unknown.

There are two main mechanisms of fat mass expansion: hypertrophy (increase in size of mature adipocytes) and hyperplasia (increase in number of mature adipocytes). Estrogen has been implicated to play a role in both of these processes. Estrogens regulate hypertrophy by modulating lipolysis and lipogenesis of mature adipocytes (Gavin, et al. 2013; Gormsen, et al. 2012; Monjo, et al. 2005; Stubbins et al. 2012; Zang, et al. 2007), and hyperplasia via affecting the differentiation of adipocyte precursors (APs) (Jeffery, et al. 2016). As mature adipocytes are post-mitotic, the generation of new adipocytes requires the proliferation and differentiation of APs (Berry and Rodeheffer 2013; Cristancho and Lazar 2011; Rodeheffer, et al. 2008). Recent work from our lab has elucidated the role of sex differences in adipocyte hyperplasia (Jeffery, et al. 2015; Jeffery et al. 2016; Sebo and Rodeheffer 2021). Male mice undergo adipocyte hyperplasia in obesity specifically in visceral fat (VWAT), while female mice display hyperplasia in both VWAT and SWAT. However, OVX females have VWAT-specific AP proliferation on HFD, similar to males (Jeffery et al. 2016). Furthermore, treating males with estrogen when on a HFD increases SWAT AP proliferation (Jeffery et al. 2016). Therefore, estrogen appears to influence the sexual dimorphic distribution of WAT in obesity via adipocyte hyperplasia in SWAT, but the cellular and molecular mechanisms are not understood. In this study, we use targeted models of ERα deletion to study the role of estrogen signaling in the adipocyte cellular lineage in obesity.

## Materials and Methods

### Animals

The Institutional Animal Care and Use Committee (IACUC) at Yale University approved all animal studies. All animals were kept in temperature and humidity-controlled rooms on a 12-hour:12-hour light:dark cycle, with lights on from 7:00 a.m. to 7:00 p.m. All mice used for these studies were on the C57BL/6J genetic background. *Adipoq-cre* mice (stock no. 028020) and *PdgfRα-cre* mice (stock no. 013148) were purchased from Jackson Laboratories. Esr1^fl^ mice were a generous gift from Dr. Sean Morrison (UT Southwestern, Dallas, TX, USA). All cre lines were crossed to the mTmG (stock no. 007676) mice purchased from Jackson Laboratories. Breeding was done in the Yale Animal Resource Center and mice were weaned at p21. Unless otherwise noted, mice were males and females 6-9 weeks of age at the start of experiments. VWAT refers to the perigonadal WAT and SWAT refers to the inguinal WAT in mice. Body composition measures were done with NMR using the Echo MRI whole body composition analyzer (Echo Medical System, Houston, TX). High-fat diet is from Research Diets (D12492). Standard diet is from Harlan Laboratories (2018S).

### RNA extraction and cDNA synthesis

For gene expression analysis, whole tissues were collected and stored at −80C until RNA extraction was performed. For adipocyte fractionation, WAT was collected and digested with 0.8mg/mL collagenase type 2 for 45 – 60 min in a shaking water bath at 37C. Samples were passed through 40um filter and centrifuged 300 g for 3 minutes. Stromal vascular fraction (SVF) was removed using oral gavage needle and adipocyte layer was left intact. Both SVF and adipocytes were rinsed with DPBS (Life Technologies) and resuspended in Trizol reagent. RNA was isolated using Direct-zol RNA Miniprep Kit (Zymo Research), according to manufacturers’ instructions. RNA was quantified by a nanodrop spectrophotometer (Thermo Fisher Scientific) and single stranded cDNA was synthesized from total RNA using the High-Capacity cDNA reverse Transcription Kit (Applied Biosystems, Life Technologies) according to the manufacturer’s instructions.

### Quantitative real-time PCR

Quantitative RT-PCR (qPCR) was performed on the cDNA using the Sybr green method of quantification on a Roche Lightcycler 480 using a SYBR FAST quantitative PCR kit (Kapa Biosystems; KK4611). Gene expression was analyzed for each sample in triplicate using the forward and reverse primers indicated on Table 1. Housekeeping genes were tbk1 or b-actin as deemed appropriate. For each experiment, cDNA samples were pooled and a standard curve was generated to quantify relative mRNA transcript levels. To confirm effective adipocyte fractionation, expression of *adipoq* and *pdgfRα* were assessed in SVF and adipocyte fractions.

**Table 1.**
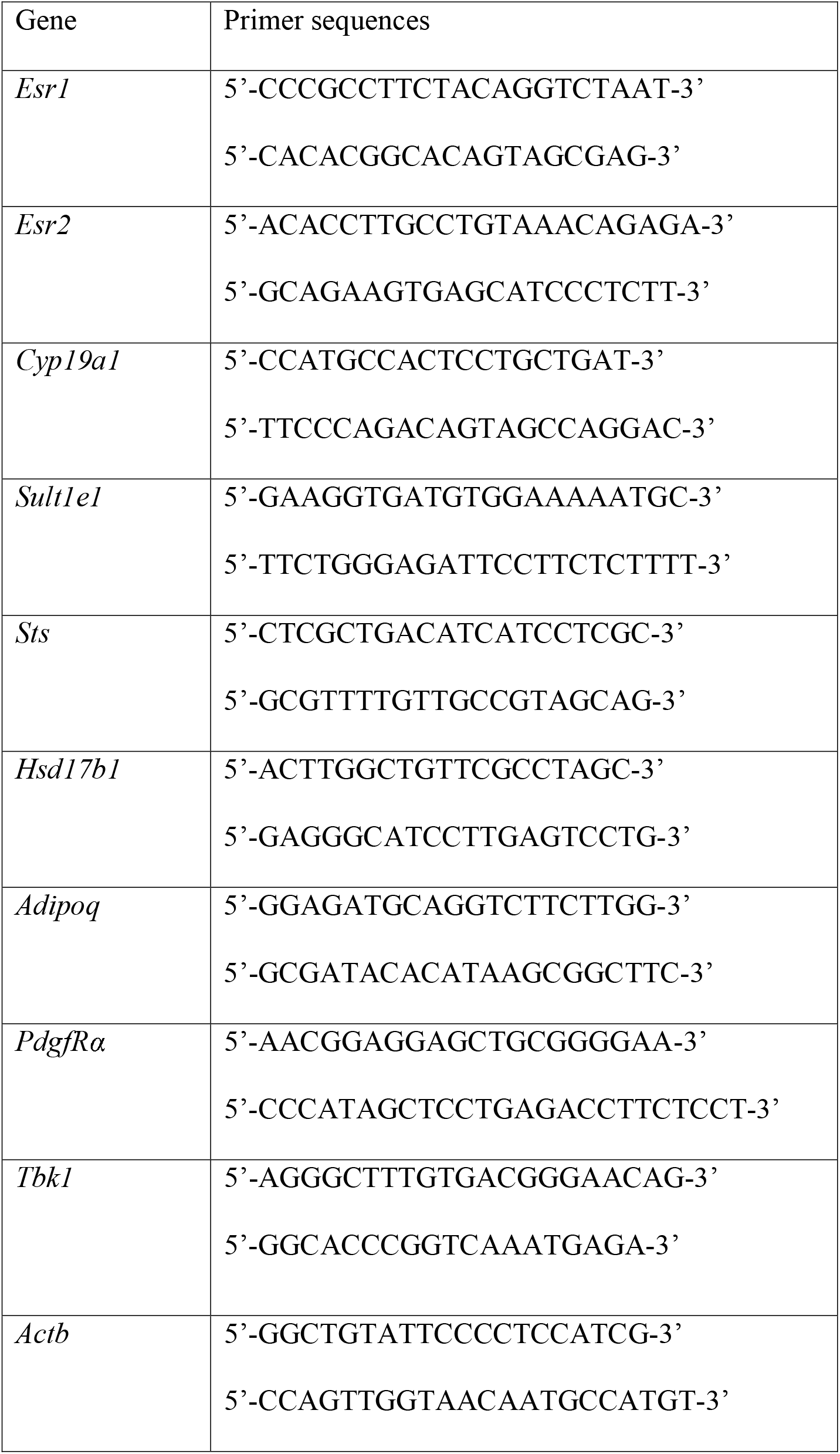
Primer sequences for gene expression analysis

### BrdU treatments

For BrdU experiments, BrdU (US Biological, B2850) was given in the drinking water at 0.8 mg/mL for experiments lasting one week or less and 0.4 mg/mL for experiments lasting more than one week. BrdU water was replenished every 48 hours.

### Confocal microscopy

Adipose tissue was collected, embedded in paraffin, and stained as previously described (Holtrup, et al. 2017; Jeffery et al. 2015; Jeffery et al. 2016; Sebo and Rodeheffer 2021). For adipocyte nuclei analysis, 20-30 images for every tissue section were acquired at 40X with a Leica TCS SP5 confocal microscope. Quantification of BrdU in adipocyte nuclei was done as previously described (Jeffery et al. 2015). At least 50 adipocyte nuclei were scored for each animal.

For adipocyte diameter measurements, the area of each adipocyte (in square pixels) was measured using Cell Profiler. The diameter of each adipocyte was calculated using the measured area, assuming each adipocyte is a perfect circle. At least 200 adipocytes were measured for each animal.

For whole mount microscopy, tissues were dissected and cut into ~1.5×1.5 cm pieces. Samples were subsequently mounted onto microscope slides with Fluoromount-G (SouthernBiotech, 0100-01) and imaged at 20X with a Leica TCS SP5 confocal microscope.

### Flow cytometry

Flow cytometry was performed as described previously (Jeffery et al. 2015) for BrdU analysis with the following antibodies: CD45 APC-eFluor 780 (eBioscience; 47–0451–80) at 1:1,000, CD31 PE-Cy7 (eBioscience, 25–0311–82), at 1:500, CD29 Alexa Fluor 700 (BioLegend, 102218) at 1:400 and Sca-1 V500 (BD Horizon, 561228) at 1:300. Cells were washed and then fixed and permeabilized using Phosflow lyse/fix and Perm Buffer III (BD Biosciences) according to the manufacturer’s recommendations. Cells were then treated with DNase (deoxyribonuclease I; Worthington; x 200 units/ml) in DPBS (Sigma; with calcium chloride and magnesium chloride) for 2-hrs at 37C and then washed in HBSS with 3% BSA. Cells were then stained with anti-BrdU antibody (Alexa Fluor 647; Phoenix Flow Systems; AX647) at 1:30 in HBSS with 3% BSA overnight in the dark at 4C. Cells were then washed in HBSS with 3% BSA and incubated with CD34 Brilliant Violet 421 (BioLegend 230 119321) at 1:400 and CD24 PerCP-Cyanine 5.5 (eBioscience, 45–0242–80) at 1:250. Following antibody incubation, samples were washed and analyzed on a BD LSRII analyzer. Data analysis was performed using BD FACS Diva software (BD Biosciences).

### Hormone quantification

Plasma 17-β estradiol was measured using Cayman’s ELISA assay (No. 501890). All other sex hormones were measured with LC-MS/MS by OpAns LLC.

### Transplant Assay

For cell transplant assays, APs were isolated and transplanted in WAT as previously described (Jeffery et al. 2016; Rodeheffer et al. 2008). For mTmG samples, NucRed™ Live 647 ReadyProbes™ Reagent (Thermo Fischer, R37106) was used as live/dead stain at 1:600 dilution. VWAT and SWAT of AP-ERαKO animals were pooled and GFP expression confirmed via FACS. Recipient male C57Bl/6J mice were anesthetized with isoflurane and surgeries performed using sterile technique. 0.5-1 million ERαKO AP cells were re-suspended in 15uL of PBS and injected into the left SWAT of 4–5-week-old congenic wildtype mice. A control PBS injection was done in the right SWAT. Mice were allowed to recover for 2 weeks, then placed on a HFD and treated with BrdU for 1 week. Left and right SWAT tissues were collected and analyzed separately via flow cytometry for incorporation with BrdU. ERαKO APs were identified by GFP fluorescence. Results were counted only for transplants in which more than 100 individual GFP-positive donor AP cells were recovered in the recipient SWAT.

### Statistical Analysis

Statistical analyses are described in each figure legend. All tests were performed using GraphPad Prism version 9.0. Data are presented as mean ± s.e.m. and p<0.05 was considered statistically significant. A minimum of 5 animals were used for each experiment, unless statistical significance was reached with fewer animals. Sample size is indicated in each figure legend. Experiments were not blinded, as genotypes of mice were known prior to analysis.

## Results

### Adipocyte Estrogen Receptor alpha (ERα) signaling does not contribute to obesity

Estrogen signaling plays an important role in both female and male physiology, as both sexes of whole-body ERα-KO and Aromatase-KO mice have significant WAT accumulation via both hypertrophy and hyperplasia (Heine et al. 2000; Jones et al. 2000; Jones et al. 2001). To determine if estrogen affects mature adipocyte function directly, we crossed adipocyte-specific *Adiponectin-cre* mice (Eguchi, et al. 2011) to *Esr1^fl/fl^* mice (Feng, et al. 2007) to create Adi-ERα-KO mice (Figure 1A). To confirm the knockout model is effective, we isolated mature adipocytes and stromal vascular fraction (SVF) in WAT and performed quantitative PCR. After confirming effective adipocyte isolation by enrichment of *adipoq* (Supp. Figure 1A, 1C), we found a 90% reduction in ERα transcripts in adipocytes from both VWAT and SWAT in males and females (Figure 1B-C) with no impact on ERα expression in SVF (Supp. Figure 1B, 1D), demonstrating efficient and specific knockout of ERα. To determine if loss of ERα affects WAT accumulation, we fed Adi-ERα-KO mice and control littermates (ERα f/f) standard diet (SD) or HFD for 8 weeks and measured body composition via echo-MRI throughout. Adi-ERα-KO females had no differences in body weight accumulation in either diet (Figure 1D-E) but had a reduction in total fat mass after a HFD (Supp. Figure 2A-B). Adi-ERα-KO males on a HFD do not show differences in body composition (Figure 1F-G, Supp. Figure 3A-B), but Adi-ERα-KOs are slightly heavier than controls on a SD (Figure 1F). This increase is not due to differences in fat mass or lean mass (Figure 1G, Supp. Figure 3A-B). Together, these data suggest that adipocyte ERα signaling does not contribute to the obesity phenotypes seen in whole body ERα-KO males and females.

**Figure 1.**
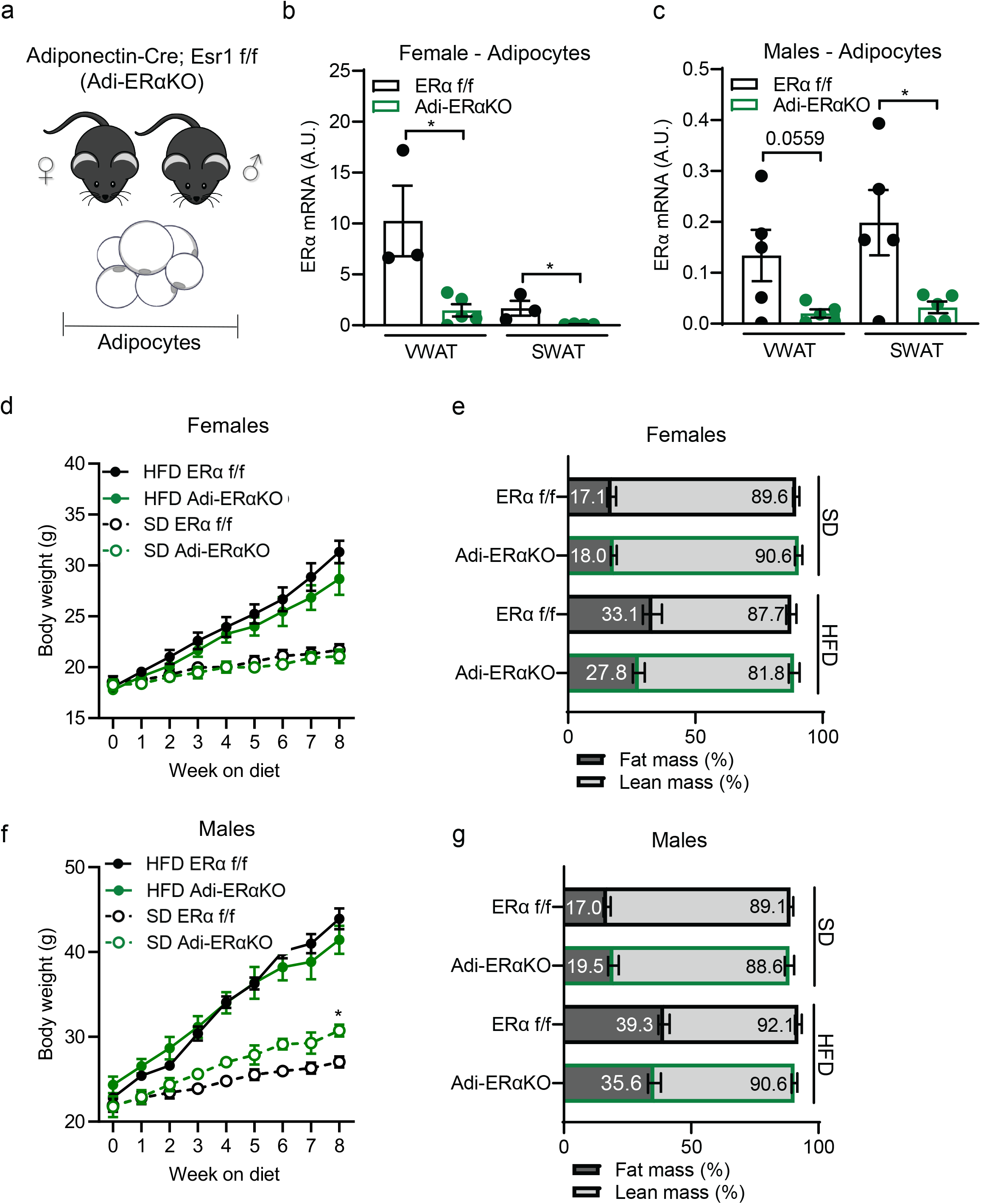
Adipocyte ERα deletion does not impact body composition. **(A)** Experimental mouse model to delete ERα in mature adipocytes using the *Adiponectin-cre*; *Esr1 fl/fl;mTmG* strain. **(B)** ERα expression in isolated mature adipocytes from Adi-ERα-KO females and controls. (n=3-5 mice per group) **(C)** ERα expression in isolated mature adipocytes from Adi-ERα-KO males and controls. (n=5 mice per group) **(D)** Body weight of female Adi-ERα-KO and controls during 8 weeks of SD or HFD. (n= 6-7 mice per group) **(E)** Body composition of female Adi-ERα-KO and controls after 8 weeks of SD and HFD. (n=6-7 mice per group) **(F)** Body weight of male Adi-ERα-KO and controls during 8 weeks of SD and HFD. Comparison shown is between SD groups. (n= 5-6 mice per group) **(G)** Body composition of male Adi-ERα-KO and controls after 8 weeks of SD and HFD. (n=5-6 mice per group) Statistical significance determined by unpaired t-tests in panel B-C and two-way ANOVA with Tukey’s test for panels C and E. *p<0.05. Error bars represent mean ± S.E.M. SD: standard diet, HFD: high-fat diet, VWAT: visceral fat, SWAT: subcutaneous fat. See also Figure S1.

### Adipocyte ERα signaling modestly impacts WAT accumulation in female mice

Even though there are no differences in body composition overall in Adi-ERα-KO mice, we assessed if differences in WAT distribution were present. After 8 weeks of HFD, both Adi-ERα-KO and control females accumulate more WAT than females on SD (Figure 2A). However, we found a significant reduction in VWAT mass in Adi-ERα-KO females on a HFD and a trend towards a reduction in SWAT (Figure 2A). Of note, there were no significant differences between liver or intrascapular brown adipose (iBAT) weights (Supp Figure 2C-D). In male Adi-ERα-KO mice, there are no differences in VWAT or SWAT accumulation on either diet (Supp. Figure 3C) or liver and iBAT weights (Supp. Figure 3D-E). These data indicate that adipocyte ERα does not play a role in male WAT expansion and only a minor role in female WAT expansion.

**Figure 2.**
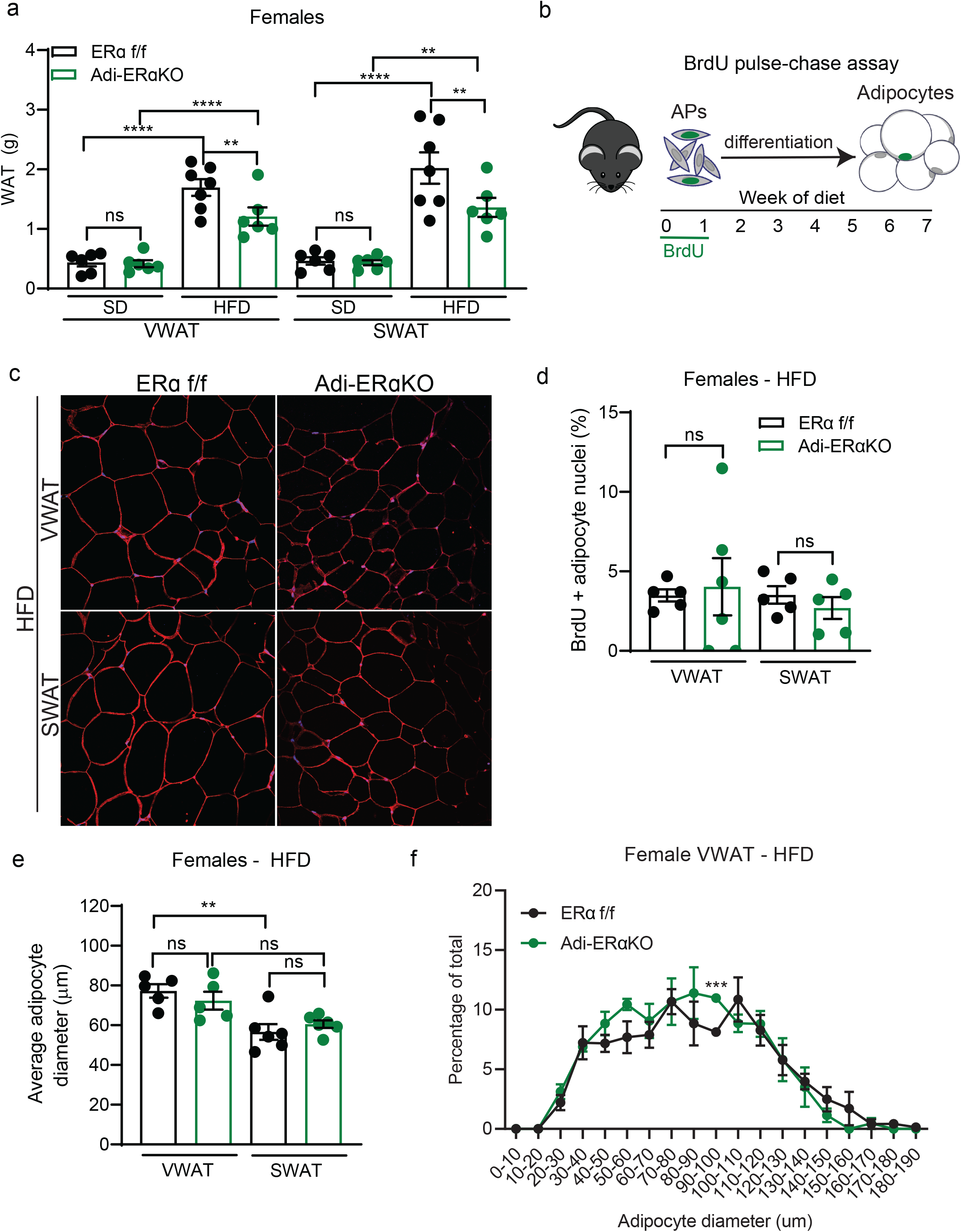
Adipocyte ERα deletion modestly impacts VWAT accumulation in HFD-fed female mice. **(A)** VWAT and SWAT accumulation of Adi-ERα-KO and control females after 8 weeks of diet. (n=6-7 mice per group) **(B)** Experimental design for BrdU pulse-chase to measure adipocyte formation. **(C)** Representative confocal microscopy images of WAT from female Adi-ERα-KO and controls after 8 weeks of HFD. Tissue stained for caveolin (red), BrdU (green), and DAPI (blue) taken at 40X. **(D)** Quantification of BrdU in adipocyte nuclei of female Adi-ERα-KO mice and controls after 8 weeks of HFD. (n=5-6 mice per group) **(E)** Average adipocyte diameter (μm) of female Adi-ERα-KO and controls after 8 weeks of HFD. (n=5-6 per group) **(F)** Distribution of adipocyte diameters in VWAT of female Adi-ERα-KO and controls after 8 weeks of HFD. (n=5-6 per group) Statistical significance determined by two-way ANOVA with Tukey’s test for panels A, D, and E. Statistical significance determined by multiple unpaired t-tests for panel F. Error bars represent mean ± S.E.M. **p<0.01, ***p<0.001, ****p<0.0001. SD: standard diet, HFD: high-fat diet, VWAT: visceral fat, SWAT: subcutaneous fat, AP: adipocyte precursors. See also Figure S2-S3.

To determine if the reduction of VWAT in Adi-ERα-KO females on a HFD is due to changes in adipocyte hyperplasia, we employed a BrdU pulse-chase assay (Figure 2B) to measure the formation of new adipocytes, as described previously (Jeffery et al. 2015; Jeffery et al. 2016). After staining with caveolin to label the adipocyte plasma membrane, BrdU, and DAPI (Figure 2C), we found no differences in BrdU incorporation into mature adipocyte nuclei of female Adi-ERα-KO mice on a HFD (Figure 2D). To determine if adipocyte hypertrophy is affected in this model, we measured the diameter of adipocytes in WAT. Although there are no differences on the average adipocyte size between the HFD groups (Figure 2E), we found a slight VWAT-specific shift towards smaller adipocytes in Adi-ERα-KO females (Figure 2F, Supp. Figure 2E). Of note, VWAT adipocytes were larger than SWAT adipocytes in both groups (Figure 2E). These data indicate that lack of ERα signaling in adipocytes has a small effect on adipocyte size an does not affect adipocyte formation in female mice on HFD. Overall, the findings of the hyperplasia and hypertrophy assays indicate adipocyte-intrinsic ERα signaling does not significantly impact fat mass accumulation, even in obesity, in males or females.

### Ablation of ERα in adipocyte precursors changes obesogenic AP proliferation patterns in mice

As loss of ERα in adipocytes does not significantly impact obesity in mice, we next focused on a potential role of ERα in APs in obesity. These cells do not express *adipoq* and therefore are not targeted in the Adi-ERα-KO mouse model (Berry and Rodeheffer 2013; Lee, et al. 2012). Thus, we deleted ERα in APs by crossing the *PdgfRα-cre* strain (Figure 3A) (Berry and Rodeheffer 2013) to *Esr1^fl/fl^* mice (Feng et al. 2007) to create the AP-ERα-KO mice. It is important to note that the mature adipocytes in this model also lack ERα. When quantifying the expression of ERα, we found the mice have a significant reduction of ERα in WAT from both male and female mice (Figure 3B). Furthermore, ERβ remains unchanged and lowly expressed in WAT (Supp. Figure 4A). To test obesogenic AP proliferation, we used flow-cytometry to measure BrdU incorporation into AP cells after 1 week of SD or HFD (Berry and Rodeheffer 2013; Jeffery et al. 2015; Jeffery et al. 2016; Rodeheffer et al. 2008). In this assay, we found female AP-ERα-KO still have HFD-induced AP proliferation, but, the APs in the AP-ERα-KO females have a more consistent proliferative response to HFD that is similar to the maximal response observed in control female mice (Figure 3C). Male AP-ERα-KO mice display normal HFD-induced VWAT AP proliferation, however, unlike controls, these male mice also have increased SWAT AP proliferation on a HFD (Figure 3D). Thus, male AP-ERα-KO have a female pattern of HFD-induced AP proliferation. These data indicate that deleting ERα with *PdgfRα-cre* affects AP proliferation in male mice upon HFD feeding.

**Figure 3:**
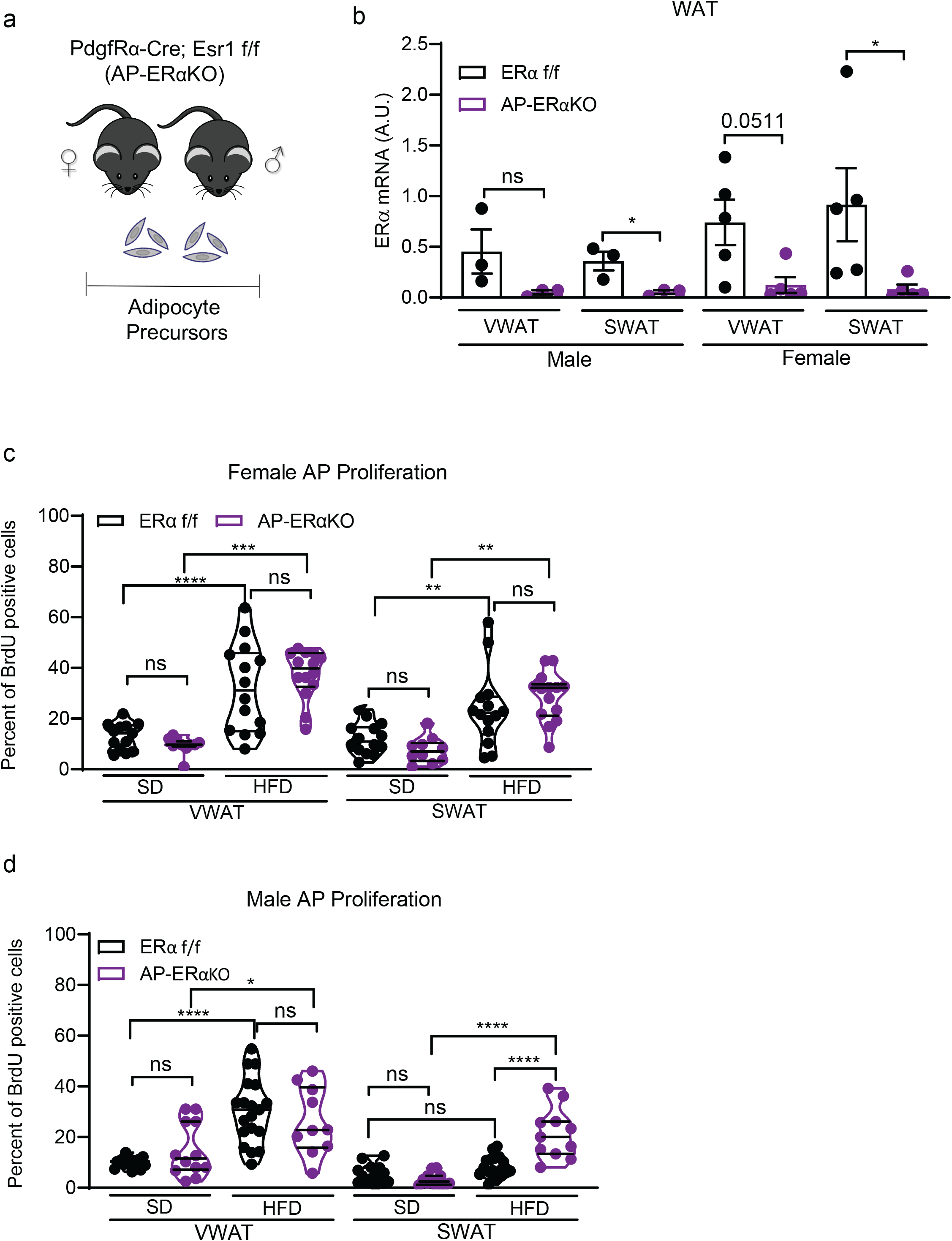
Lack of ERα in adipocyte precursors (APs) changes obesogenic proliferation patterns. **(A)** Experimental mouse model to delete ERα in APs using *PdgfRα-cre*; *Esr1 fl/fl*;*mTmG* strain. **(B)** ERα expression in whole white adipose tissue (WAT) from AP-ERα-KOs and controls. (n=3-5 mice per group) **(C)** AP proliferation of female AP-ERα-KOs and controls after one week of SD or HFD. (n=10-14 mice per group) **(D)** AP proliferation of male AP-ERα-KOs and controls after one week of SD or HFD. (n=10-19 mice per group) Statistical significance determined by unpaired t-tests for panel B. Statistical significance determined by two-way ANOVA with Tukey’s test for panels C-D. In dot plots, error bars represent mean ± S.E.M. In violin plots, mean is showed as filled line and quartiles as dotted lines. *p<0.05, **p<0.01, ***p<0.001, ****p<0.0001. SD: standard diet, HFD: high-fat diet, VWAT: visceral fat, SWAT: subcutaneous fat, AP: adipocyte precursors. See also Figure S4.

### Ablation of ERα in adipocyte precursors causes increased fat mass accumulation upon HFD feeding

As AP-ERα-KO females have HFD-induced AP proliferation, we next assessed if this proliferation is indeed adipogenic. AP-ERα-KO females and controls were fed a SD or HFD for 8 weeks and body composition was assessed weekly. Interestingly, AP-ERα-KO females gained significant more weight on the HFD compared to controls (Figure 4A) with increased fat and lean mass accumulation (Figure 4B). Although no changes in VWAT and SWAT mass, and thus fat distribution, were found, AP-ERα-KO females consistently gain fat mass on a HFD (Figure 4C). There are no differences in WAT accumulation on a SD (Figure 4C), indicating these phenotypes are HFD-specific.

**Figure 4:**
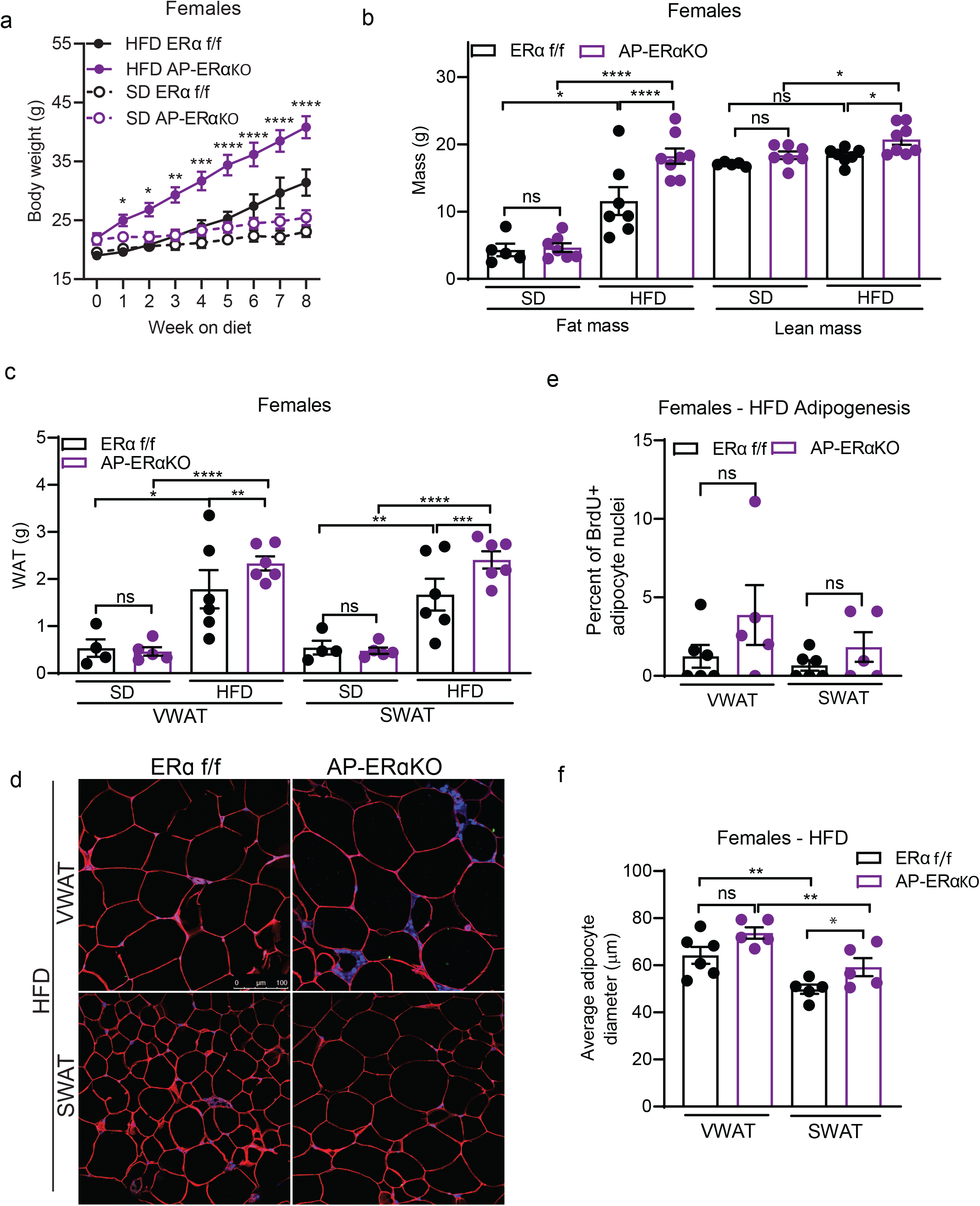
AP-ERα-KO female mice are more susceptible to diet-induced obesity. **(A)** Body weight of female AP-ERα-KOs and controls during 8 weeks of SD and HFD. Significance shown between HFD groups. (n=5-8 mice per group) **(B)** Total fat and lean mass of female AP-ERα-KOs and controls after 8 weeks of SD and HFD. (n=5-8 mice per group) **(C)** VWAT and SWAT accumulation of female AP-ERα-KOs and controls after 8 weeks of SD and HFD. (n=5-8 mice per group) **(D)** Representative confocal microscopy images of WAT from female AP-ERα-KO and controls after 8 weeks of HFD. Tissue stained for caveolin (red), BrdU (green), and DAPI (blue) taken at 40X. Scale bar is 100μm. **(E)** Quantification of BrdU in adipocyte nuclei of female AP-ERα-KO mice and controls after 8 weeks of HFD. (n=5-6 mice per group) **(F)** Average adipocyte diameter (μm) of female AP-ERα-KO and controls after 8 weeks of HFD. (n=5-6 per group) Statistical significance determined by two-way ANOVA with Tukey’s test. Error bars represent mean ± S.E.M. *p<0.05, **p<0.01, ***p<0.001, ****p<0.0001. SD: standard diet, HFD: high-fat diet, VWAT: visceral fat, SWAT: subcutaneous fat. See also Figure S5.

Next, we determined if the differences in fat mass in AP-ERα-KOs were caused by increased hypertrophy or hyperplasia using the BrdU pulse-chase assay (figure 2B) to quantify the formation of newly formed adipocytes (Figure 4D). Interestingly, we find a trend towards more hyperplasia in VWAT and SWAT of AP-ERα-KO females on HFD (Figure 4E). When adipocyte size was measured, AP-ERα-KO females on a HFD showed a trend towards larger adipocytes in both VWAT and SWAT, but this result did not reach statistical significance (Figure 4F). The weights of liver and iBAT were not altered in AP-ERα-KO females (Supp. Figure 5A-B). These data indicate that deleting ERα using *PdgfRα-cre* promotes obesity in female mice by affecting both adipocyte hyperplasia and hypertrophy.

As male AP-ERα-KO mice exhibited increased AP proliferation in SWAT, we determined if this led to long-term obesity with increased SWAT expansion. We fed AP-ERα-KO males SD or HFD for 8 weeks and found a significant increase in body weight on HFD, but not SD (Figure 5A). Similar to female AP-ERα-KO, this is the result of an increase in both fat mass and lean mass accumulation (Figure 5B). In line with increased SWAT AP proliferation, AP-ERα-KO males accumulate more SWAT on a HFD (Figure 5C). When we measured hyperplasia using the BrdU pulse-chase assay (Figure 5D), we found male AP-ERα-KO trend towards more adipogenesis in VWAT (Figure 5E) and significant increase in obesogenic hyperplasia in SWAT (Figure 5F). This is in contrast to control males where there is no hyperplasia in SWAT (Figure 5F), as has been previously reported (Jeffery et al. 2015). Furthermore, the size of SWAT adipocytes on a HFD is not changed in AP-ERα-KO males, suggesting the increase in SWAT mass is driven by adipocyte hyperplasia specifically (Supp. Figure 6A).

**Figure 5:**
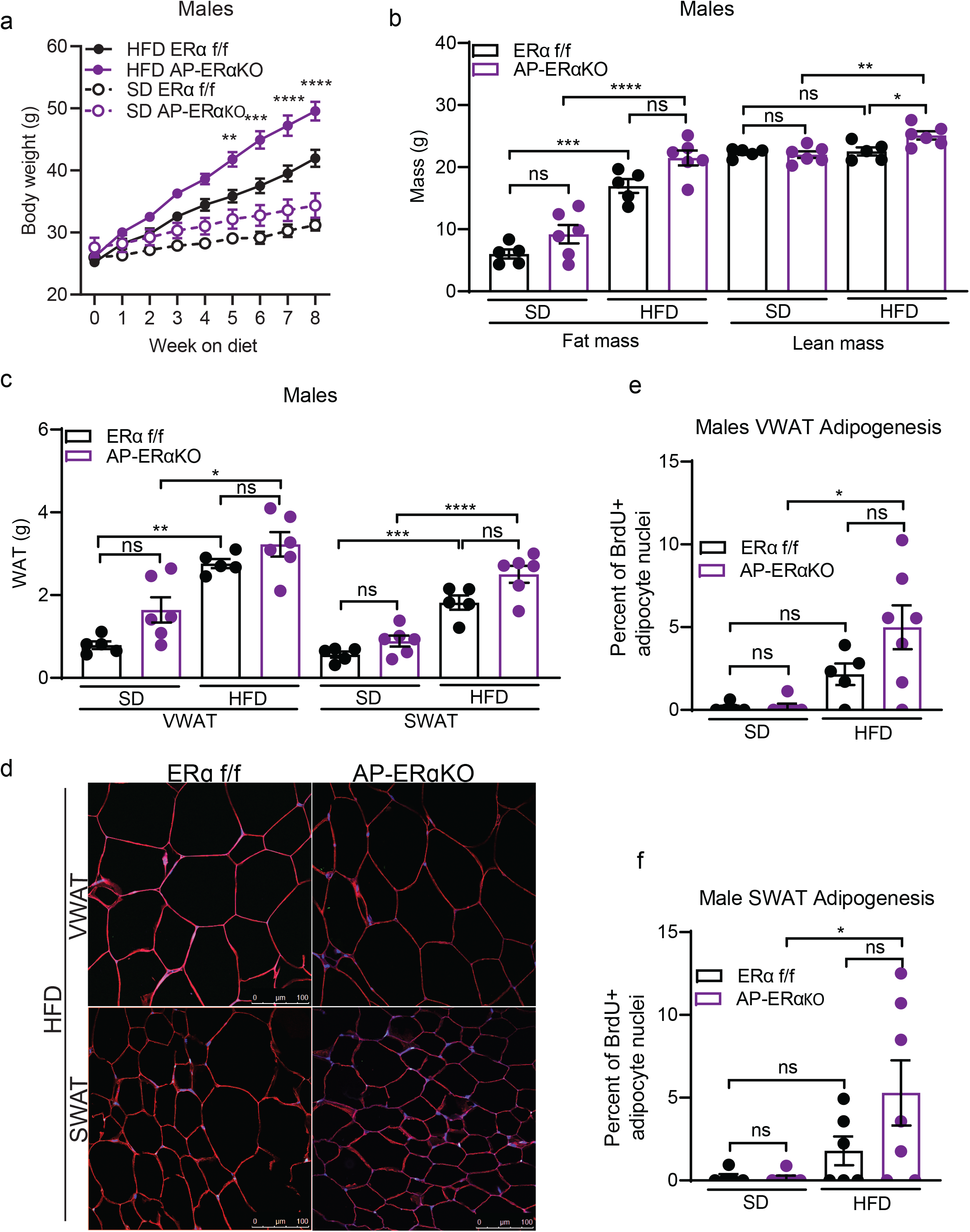
AP-ERα-KO male mice have diet-induced SWAT hyperplasia. **(A)** Body weight of male AP-ERα-KOs and controls during 8 weeks of SD and HFD. Significance shown between HFD groups. (n=5-6 mice per group) **(B)** Total fat and lean mass of male AP-ERα-KOs and controls after 8 weeks of SD and HFD. (n=5-6 mice per group) **(C)** VWAT and SWAT accumulation of male AP-ERα-KOs and controls after 8 weeks of SD and HFD. (n=5-6 mice per group) **(D)** Representative confocal microscopy images of WAT from male AP-ERα-KOs and controls after 8 weeks of HFD. Tissue stained for caveolin (red), BrdU (green), and DAPI (blue) taken at 40X. Scale bar is 100μm. **(E)** Quantification of BrdU in adipocyte nuclei in VWAT of male AP-ERα-KOs and controls after 8 weeks of SD and HFD. (n=5-6 mice per group) **(F)** Quantification of BrdU in adipocyte nuclei in SWAT of male AP-ERα-KOs and controls after 8 weeks of SD and HFD. (n=5-6 mice per group) Statistical significance determined by two-way ANOVA with Tukey’s test. Error bars represent mean ± S.E.M. *p<0.05, **p<0.01, ***p<0.001, ****p<0.0001. SD: standard diet, HFD: high-fat diet, VWAT: visceral fat, SWAT: subcutaneous fat. See also Figure S6.

While there are overt phenotypes in the SWAT of male AP-ERα-KO mice, changes in VWAT accumulation were more subtle. These mice maintained the obesogenic VWAT expansion and adipocyte hyperplasia observed in control mice fed a HFD (Figure 5C, 5E). On a SD, AP-ERα-KO males have increased VWAT mass compared to controls (Figure 5C). Interestingly, this is due to larger adipocytes and not an increase in adipocyte hyperplasia (Figure 5E, Supp. Figure 6A). In contrast, no significant changes in mass, size, or hyperplasia were seen in SWAT of AP-ERα-KO males on SD (Figure 5C, 5E-F, Supp. Figure 6A). Of note, no changes in iBAT mass were observed (Supp. Figure 6B), but AP-ERα-KO males had larger livers on a HFD compared to controls (Supp. Figure 6C). Together, these data demonstrate that AP-ERα-KO males on a HFD have a more feminized pattern of obesity, displaying increased AP proliferation and hyperplasia in SWAT.

### AP-ERα-KO mice have altered sex hormone levels

Many mouse models of hormone receptor knockouts or inhibition via drug treatments result in excessive production of sex hormones, with several ERα knockout models having increased estrogen in plasma (Curtis Hewitt, et al. 2000; Gustafsson, et al. 2016; Kauffman et al. 2015; Xu, et al. 2011). Thus, we measured 17-β estradiol in plasma of AP-ERα-KO mice and found significantly higher levels in female KOs compared to controls (Figure 6A). In line with this finding, the expression of enzymes that produce and metabolize estrogens are significantly enriched in the gonads of AP-ERα-KO female mice (Figure 6B). Interestingly, AP-ERα-KO females also have elevated testosterone and androstenedione levels (Figure 6C, Supp. Figure 7A). Although to a lesser extent than females, males are capable of producing estrogen in the testis, brain, skin, and adipose tissue (Baird, et al. 1973; Liu, et al. 2013; MacDonald, et al. 1979; Ohlsson, et al. 2017; Schneider, et al. 1979). However, when we measured 17-β estradiol in male plasma, all but one sample fell below the detection limit (Supp Figure 7B), however, we do find increased testosterone and androstenedione in the plasma of AP-ERα-KO males (Figure 6C, Supp. Figure 7A). These data suggest that AP-ERα-KO mice have compensatory mechanisms that lead to elevated levels of multiple sex hormones, thus making it a difficult model to assess the cellular mechanism of ERα in APs as both androgens and estrogens have been implicated to play a role in hyperplasia and fat mass regulation (Jeffery et al. 2016; Sebo and Rodeheffer 2021).

**Figure 6:**
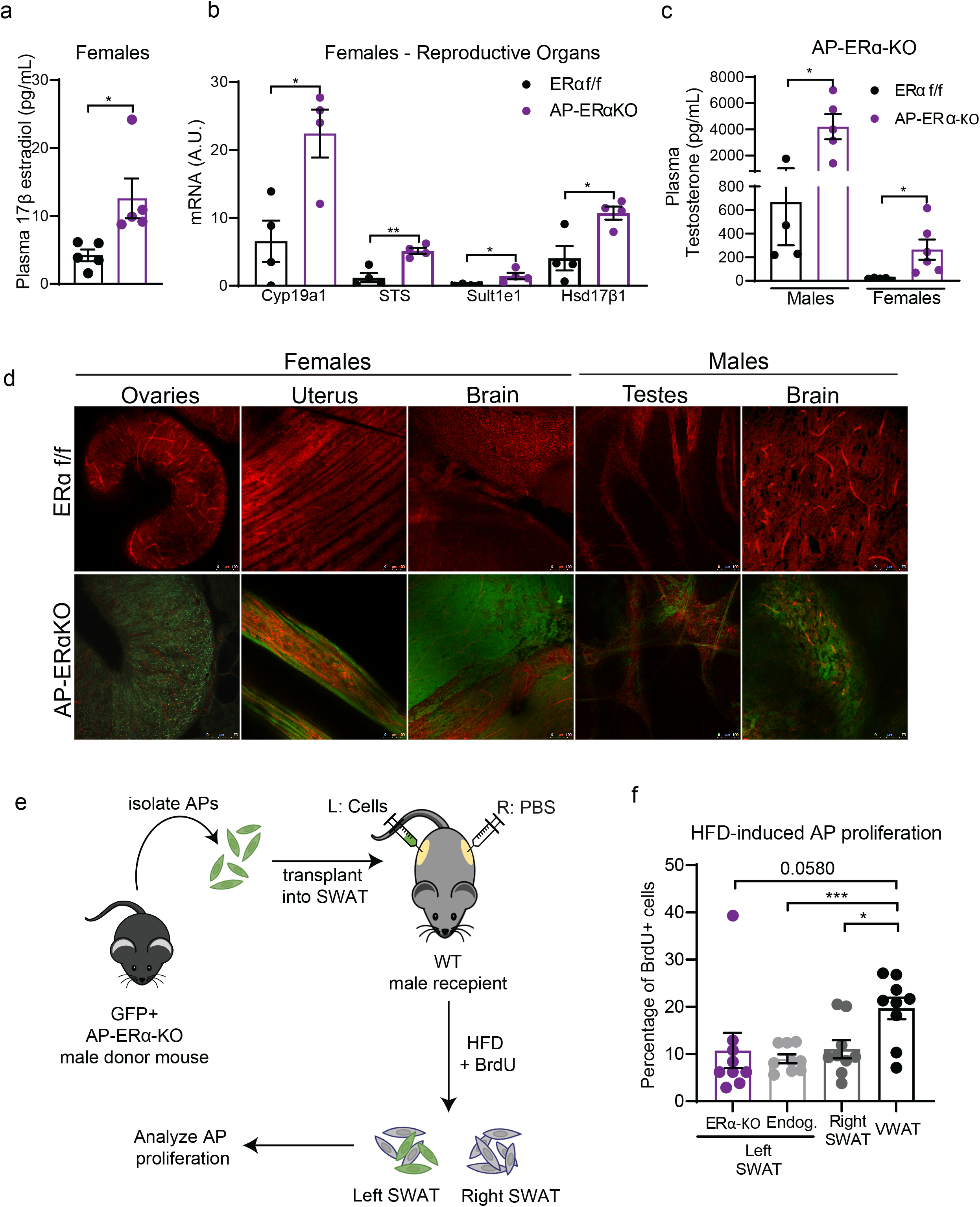
Off-target effects in AP-ERα-KO mice drive obesogenic hyperplasia. **(A)** Plasma levels of 17-β estradiol in female AP-ERα-KOs and controls. (n=5 mice per group) **(B)** Expression of estrogen metabolizing genes in the reproductive organs of AP-ERα-KO and control females. (n=4 mice per group) **(C)** Plasma levels of testosterone in male and female AP-ERα-KOs and controls. (n=4-6 per group) **(D)** Whole-mount confocal images of tissues from *AP-ERα-KO; mTmG* mice taken at 20X. GFP fluorescence indicates presence of cre activity. Scale bar is 100μm. (n=1 mice per group) **(E)** Experimental design of AP transplant experiment. AP cells lacking ERα were injected into the SWAT of wildtype C57BL/6J male mice. AP proliferation was measured after one week of HFD. **(F)** One week AP proliferation of transplanted ERα-KO cells, endogenous cells from left SWAT, right SWAT (PBS injection), and VWAT of recipient males. (n=9 mice per group) Statistical significance determined by unpaired t-tests. Error bars represent mean ± S.E.M. * p<0.05, **p<0.01, ***p<0.001. HFD: high-fat diet, VWAT: visceral fat, SWAT: subcutaneous fat. See also Figure S7-9.

Although *PdgfRα-cre* effectively targets the adipocyte lineage in adipose tissue (Berry and Rodeheffer 2013), its expression is not restricted to APs or WAT (Roesch, et al. 2008). Thus, we assessed other sites where ERα expression may be affected in the AP-ERα-KO mice using the dual fluorescent reporter mTmG (Muzumdar, et al. 2007). Upon expression of cre recombinase, cells switch from expressing plasma membrane-targeted Tomato (red fluorescence) to expressing plasma membrane-targeted GFP. Therefore, GFP fluorescence can be used to trace cre recombinase activity. Next, we extensively characterized WAT as well as other tissues from male and female AP-ERα-KO mice using whole-mount confocal microscopy as done previously (Berry and Rodeheffer 2013; Jeffery, et al. 2014; Sebo, et al. 2018). As anticipated, we found GFP expression from VWAT, SWAT, and iBAT but not in liver or muscle (Supp. Figure 8A). When other tissues were imaged, we found GFP expression in the brain and gonadal tissues of both male and female mice (Figure 6D). While it has been published that a population of glial cells express PdgfRα (Roesch et al. 2008), we did not anticipate to see GFP expression in the gonads of males and females.

Due to the important role of estrogens in the female reproductive system, we looked further into the physiological effects of ERα loss via *PdgfRα-cre* on the gonads. Gross morphology of the reproductive tract showed atrophied uterus and hemorrhagic ovaries, consistent with previous models of ERα deletion (Figure 6D, Supp. Figure 9A-B) (Antonson, et al. 2014; Lubahn, et al. 1993). Next, we confirmed reduced ERα transcripts by qPCR in the female gonads (Supp. Figure 9C). Of note, we found no changes in ERβ transcripts. Based on these results, we assessed if there were disturbances in the estrus cycle of AP-ERα-KO females. Interestingly, AP-ERα-KO females do not cycle and are infertile (Supp. Figure 9D-E) similar to whole-body ERα-KOs (Heine et al. 2000).

### Increased SWAT hyperplasia in AP-ERα-KO males is due to AP-extrinsic mechanisms

Our lab has previously shown that microenvironment changes in adipose tissue influence the response of APs to HFD (Jeffery et al. 2016). Adipose tissue is a major source of estrogen production in humans (Hetemäki, et al. 2017; Nelson and Bulun 2001). There is some evidence that mice adipose can produce estrogen (Chow, et al. 2009; Polari, et al. 2015; Zhao, et al. 2009), although some groups report undetectable levels in mice WAT (Kim, et al. 2021). In addition, treating male mice with 17-β estradiol induces SWAT proliferation (Jeffery et al. 2016). To determine if the increased SWAT proliferation in AP-ERα-KO males is due to tissue microenvironment changes, we attempted to quantify sex hormones within the adipose tissue of these mice via several methods, including mass spectrometry, but our efforts were unsuccessful (data not shown). As an alternative approach, we employed an AP transplant assay to test if the proliferation of these ERα-KO cells was sustained in a wildtype tissue microenvironment. Briefly, we sorted GFP+ AP cells from AP-ERα-KO males and injected them into the SWAT of wildtype C57BL/6J males (Figure 6E). After 2 weeks of recovery, recipient mice were fed HFD and treated with BrdU for one week. AP proliferation was measured for endogenous cells (GFP-) and transplanted ERα-KO cells (GFP+) via flow cytometry. In contrast to the AP-ERα-KO mice, ERα-KO cells do not proliferate in the SWAT of wildtype male mice (Figure 6F). These data indicate that the increased AP proliferation, hyperplasia, and accumulation of SWAT in AP-ERα-KO male mice is driven by AP-extrinsic changes in the tissue microenvironment and not a lack of ERα function in APs.

## Discussion

The expansion and distribution of adipose tissue is sexually dimorphic. Previous work in the lab has shown that sex hormones play a role in this process by influencing hyperplasia (Jeffery et al. 2016; Sebo and Rodeheffer 2021). Our attempt to elucidate the role of estrogen signaling in APs led to an important lesson in the use of hormone receptor knockout models. *PdgfRα-cre* is known be expressed in several non-adipose tissues. Here we find that it also marks female and male reproductive organs (Figure 6D, Supp. Figure 9C). This deletion of ERα in one or several of the target tissues leads to compensatory mechanisms that elevate sex hormones in the plasma of AP-ERα-KO mice, including estradiol in females (Figure 6A-C, Supp. Figure 7A-B), making this model unsuitable to study ERα signaling in APs. Despite having increased estrogen, an obesity-protecting hormone, AP-ERα-KO mice are more obese. This finding replicates many other ERα-KO models using cre promoters that target the brain (Xu et al. 2011), and whole-body ERα-KO (Curtis Hewitt et al. 2000; Gustafsson et al. 2016) where the mice are more obese despite the increased estrogen levels. This highlights that ERα mediates many of estrogens protective roles against obesity. In addition, estrogen can still signal through ERβ and novel GPR30 receptors in these models. Thus, it is difficult to determine if the phenotypes we observe here are due to lack of ERα signaling in target cells or the result of alteration of sex hormones and their intermediates that can signal via ERα-independent mechanisms (Sebo and Rodeheffer 2021).

Nonetheless, our findings in AP-ERα-KO mice do support an important role of estrogens in obesity. The fact that AP-ERα-KO females are more likely to become obese than control females is of interest. Our lab has found that wildtype females have a large variation in hyperplasia (Jeffery et al. 2016) and obesity phenotypes overall (Hong, et al. 2009; Yang, et al. 2014). This indicates that estrogen is likely involved in the hyperplastic response of females to a HFD and this mechanism should be further explored. Perhaps even more important is the feminized pattern of obesity that we find in male AP-ERα-KO (Figure 3D, 5C). Although the increased expansion of SWAT in these mice occurs via hyperplasia and not hypertrophy (Figure 5F, Supp. Figure 6A), we find that lack of ERα in APs itself does not cause more proliferation (Figure 6E-F). This suggest that the hormonal milieu in SWAT of male AP-ERα-KOs provides the necessary cues for adipogenesis to occur in absence of ERα. Though our attempts to measure sex hormones within WAT were unsuccessful, the fact that wildtype males treated with estrogen have SWAT proliferation (Jeffery et al. 2016) suggests increased estrogen production in AP-ERα-KO is a potential mechanism to explain this phenotype, as free estradiol correlates to increased SWAT mass in men (Vermeulen, et al. 2002).

The AP-ERα-KO model also affects estrogen signaling in mature adipocytes, and although estrogens have overall protective effects against obesity, their effects in adipocyte biology are not well understood. This is the first study to address the role of adipocyte ERα in diet-induced obesity. Our data suggest that the extensive adipose tissue accumulation seen in whole body ERα-KO are not replicated in adipocyte-specific ERα-KO. Male Adi-ERα-KO mice do not show differences in fat mass accumulation (Figure 1F, Supp. Figure 3A, 3C). However, loss of adipocyte ERα in females affects VWAT expansion via a modest reduction in adipocyte size (Figure 2A, 2F) without affecting hyperplasia (Figure 2D). It is unclear if this is due to energy expenditure or food intake changes in Adi-ERα-KO females. Importantly, estrogen levels in the plasma of male and female Adi-ERα-KO mice are not different from controls. Thus, the absence of phenotypes in these mice is not explained by changes in systemic estrogen levels.

One way estrogen is believed to modulate obesity is by affecting adipocyte lipolysis and lipogenesis (Gormsen et al. 2012; Pedersen, et al. 2004). Our data demonstrates that this potential role could be mediated by ERα in female mice as some subtle differences in VWAT adipocyte sizes were present (Figure 2F). Of note, our findings in the Adi-ERα-KO mice stand in contrast to previous findings (Davis, et al. 2013). This could be due to the use of different *Adiponectin-cre* strains and their respective efficiencies as the strain used in this study has demonstrated higher recombination levels in adipocytes (Eguchi et al. 2011; Wang, et al. 2010). Overall, these data suggest that signaling of ERα in adipocytes is not a driver of the protective effects of estrogen in obesity and that there is potential VWAT-specific role of ERα in regulating adipocyte size in females. We do not know how these findings will translate to postmenopausal adipocyte-ERα signaling or obesity in aging.

Estrogens protective effects extend to other metabolic organs like the liver, as it has been shown to improve liver function and prevent hepatic steatosis. (Chow, et al. 2011; Shen and Shi 2015; Zhu, et al. 2013) The major estrogen receptor in the liver, ERα, has been shown to have oscillating activity in the liver correlating to the estrus cycle in females. (Villa, et al. 2012) This pulsatile activity during proestrus is sufficient to drive changes in gene expression to prevent lipid accumulation. This is mediated in part by ERα as liver-specific knockouts do not show these gene expression changes and both male and female liver-ERα-KOs have fatty livers. (Villa et al. 2012) In addition, OVX female mice are more prone to develop hepatic steatosis compared to sham females, but treatment with estrogen rescues this phenotype (Zhu et al. 2013) and the hepatic steatosis seen in Aromatase-KO males can be rescued by treatment with an ERα agonist. (Chow et al. 2011) Our data shows that male but not female AP-ERα-KOs had substantially larger livers on a HFD compared to controls (Supp. Figure 6C). This is not due targeting of ERα in the livers as *PdgfRα-cre* is does not label cells in the liver (Supp. Figure 8A). Interestingly, male Adi-ERα-KOs showed a trend to larger livers on a SD (Supp. Figure 3D). These data points at a potential role for estrogen in mediating organ crosstalk to promote lipid accumulation in the liver. Other possibilities are that this is driven by the distinct hormonal milieu present in AP-ERα-KO males or a product of the increased adipose tissue mass of HFD-fed AP-ERα-KO males.

Overall, these data provide important insights in the role of estrogens in diet-induced obesity. As obesity rates and its associated comorbidities continue to rise (Finkelstein, et al. 2012; Ward, et al. 2019) it is crucial to understand the mechanisms underlying this disease. Although it is evident that sex hormones are key players in the expansion of adipose, we must be cautious of conclusions derived from hormone receptor knockout mouse models as these have consistently shown to have compensatory mechanisms that elevate estrogen and other sex hormones (Curtis Hewitt et al. 2000; Gustafsson et al. 2016; Kauffman et al. 2015; Xu et al. 2011). It is important to note that the hyperplastic phenotypes seen in AP-ERα-KO mice are exclusively present in HFD conditions. Our findings suggest the hormonal milieu in AP-ERα-KO mice is not sufficient to drive hyperplastic obesity, and that a HFD is needed for this process. Identifying these nutrient signals and how estrogen and other hormones influence expansion of adipose is crucial to our understanding of the regulation of fat mass expansion in obesity.

## Author contribution statement

RSP and MSR designed experiments. RSP and NT performed experiments. RSP and NT analyzed data. RSP and MSR interpreted data and wrote the manuscript. RSP, NT, and MSR approved the final version of the (Davis et al. 2013)manuscript.

## References

Aloia JF, Vaswani A, Ma R & Flaster E 1996 Aging in women--the four-compartment model of body composition. Metabolism 45 43–48.

Andersson B, Mattsson LA, Hahn L, Marin P, Lapidus L, Holm G, Bengtsson BA & Bjorntorp P 1997 Estrogen replacement therapy decreases hyperandrogenicity and improves glucose homeostasis and plasma lipids in postmenopausal women with noninsulin-dependent diabetes mellitus. J Clin Endocrinol Metab 82 638–643.

Antonson P, Matic M, Portwood N, Kuiper RV, Bryzgalova G, Gao H, Windahl SH, Humire P, Ohlsson C, Berggren PO, et al. 2014 aP2-Cre-mediated inactivation of estrogen receptor alpha causes hydrometra. PLoS One 9 e85581.

Baird DT, Galbraith A, Fraser IS & Newsam JE 1973 The concentration of oestrone and oestradiol-17 in spermatic venous blood in man. J Endocrinol 57 285–288.

Berry R & Rodeheffer MS 2013 Characterization of the adipocyte cellular lineage in vivo. Nat Cell Biol 15 302–308.

Bilezikian JP, Morishima A, Bell J & Grumbach MM 1998 Increased bone mass as a result of estrogen therapy in a man with aromatase deficiency. N Engl J Med 339 599–603.

Carani C, Qin K, Simoni M, Faustini-Fustini M, Serpente S, Boyd J, Korach KS & Simpson ER 1997 Effect of testosterone and estradiol in a man with aromatase deficiency. N Engl J Med 337 91–95.

Chow JD, Simpson ER & Boon WC 2009 Alternative 5′-untranslated first exons of the mouse Cyp19A1 (aromatase) gene. J Steroid Biochem Mol Biol 115 115–125.

Chow JD, Jones ME, Prelle K, Simpson ER & Boon WC 2011 A selective estrogen receptor alpha agonist ameliorates hepatic steatosis in the male aromatase knockout mouse. J Endocrinol 210 323–334.

Conte FA, Grumbach MM, Ito Y, Fisher CR & Simpson ER 1994 A syndrome of female pseudohermaphrodism, hypergonadotropic hypogonadism, and multicystic ovaries associated with missense mutations in the gene encoding aromatase (P450arom). J Clin Endocrinol Metab 78 1287–1292.

Cooke PS & Naaz A 2004 Role of estrogens in adipocyte development and function. Exp Biol Med (Maywood) 229 1127–1135.

Cristancho AG & Lazar MA 2011 Forming functional fat: a growing understanding of adipocyte differentiation. Nat Rev Mol Cell Biol 12 722–734.

Curtis Hewitt S, Couse JF & Korach KS 2000 Estrogen receptor transcription and transactivation: Estrogen receptor knockout mice: what their phenotypes reveal about mechanisms of estrogen action. Breast Cancer Res 2 345–352.

Davis KE, Neinast MD, Sun K, Skiles WM, Bills JD, Zehr JA, Zeve D, Hahner LD, Cox DW, Gent LM, et al. 2013 The sexually dimorphic role of adipose and adipocyte estrogen receptors in modulating adipose tissue expansion, inflammation, and fibrosis. Mol Metab 2 227–242.

Dubuc PU 1985 Effects of estrogen on food intake, body weight, and temperature of male and female obese mice. Proc Soc Exp Biol Med 180 468–473.

Eguchi J, Wang X, Yu S, Kershaw EE, Chiu PC, Dushay J, Estall JL, Klein U, Maratos-Flier E & Rosen ED 2011 Transcriptional control of adipose lipid handling by IRF4. Cell Metab 13 249–259.

Feng Y, Manka D, Wagner KU & Khan SA 2007 Estrogen receptor-alpha expression in the mammary epithelium is required for ductal and alveolar morphogenesis in mice. Proc Natl Acad Sci U S A 104 14718–14723.

Finkelstein EA, Khavjou OA, Thompson H, Trogdon JG, Pan L, Sherry B & Dietz W 2012 Obesity and severe obesity forecasts through 2030. Am J Prev Med 42 563–570.

Gavin KM, Cooper EE, Raymer DK & Hickner RC 2013 Estradiol effects on subcutaneous adipose tissue lipolysis in premenopausal women are adipose tissue depot specific and treatment dependent. Am J Physiol Endocrinol Metab 304 E1167–1174.

Gibb FW, Homer NZ, Faqehi AM, Upreti R, Livingstone DE, McInnes KJ, Andrew R & Walker BR 2016 Aromatase Inhibition Reduces Insulin Sensitivity in Healthy Men. J Clin Endocrinol Metab 101 2040–2046.

Gormsen LC, Host C, Hjerrild BE, Pedersen SB, Nielsen S, Christiansen JS & Gravholt CH 2012 Estradiol acutely inhibits whole body lipid oxidation and attenuates lipolysis in subcutaneous adipose tissue: a randomized, placebo-controlled study in postmenopausal women. Eur J Endocrinol 167 543–551.

Gustafsson KL, Farman H, Henning P, Lionikaite V, Moverare-Skrtic S, Wu J, Ryberg H, Koskela A, Gustafsson JA, Tuukkanen J, et al. 2016 The role of membrane ERalpha signaling in bone and other major estrogen responsive tissues. Sci Rep 6 29473.

Heine PA, Taylor JA, Iwamoto GA, Lubahn DB & Cooke PS 2000 Increased adipose tissue in male and female estrogen receptor-alpha knockout mice. Proc Natl Acad Sci U S A 97 12729–12734.

Hetemäki N, Savolainen-Peltonen H, Tikkanen MJ, Wang F, Paatela H, Hämäläinen E, Turpeinen U, Haanpää M, Vihma V & Mikkola TS 2017 Estrogen Metabolism in Abdominal Subcutaneous and Visceral Adipose Tissue in Postmenopausal Women. J Clin Endocrinol Metab 102 4588–4595.

Holtrup B, Church CD, Berry R, Colman L, Jeffery E, Bober J & Rodeheffer MS 2017 Puberty is an important developmental period for the establishment of adipose tissue mass and metabolic homeostasis. Adipocyte 6 224–233.

Hong J, Stubbins RE, Smith RR, Harvey AE & Nunez NP 2009 Differential susceptibility to obesity between male, female and ovariectomized female mice. Nutr J 8 11.

Jeffery E, Church CD, Holtrup B, Colman L & Rodeheffer MS 2015 Rapid depot-specific activation of adipocyte precursor cells at the onset of obesity. Nat Cell Biol 17 376–385.

Jeffery E, Berry R, Church CD, Yu S, Shook BA, Horsley V, Rosen ED & Rodeheffer MS 2014 Characterization of Cre recombinase models for the study of adipose tissue. Adipocyte 3 206–211.

Jeffery E, Wing A, Holtrup B, Sebo Z, Kaplan JL, Saavedra-Pena R, Church CD, Colman L, Berry R & Rodeheffer MS 2016 The Adipose Tissue Microenvironment Regulates Depot-Specific Adipogenesis in Obesity. Cell Metab 24 142–150.

Jones ME, Thorburn AW, Britt KL, Hewitt KN, Wreford NG, Proietto J, Oz OK, Leury BJ, Robertson KM, Yao S, et al. 2000 Aromatase-deficient (ArKO) mice have a phenotype of increased adiposity. Proc Natl Acad Sci U S A 97 12735–12740.

Jones ME, Thorburn AW, Britt KL, Hewitt KN, Misso ML, Wreford NG, Proietto J, Oz OK, Leury BJ, Robertson KM, et al. 2001 Aromatase-deficient (ArKO) mice accumulate excess adipose tissue. J Steroid Biochem Mol Biol 79 3–9.

Kauffman AS, Thackray VG, Ryan GE, Tolson KP, Glidewell-Kenney CA, Semaan SJ, Poling MC, Iwata N, Breen KM, Duleba AJ, et al. 2015 A Novel Letrozole Model Recapitulates Both the Reproductive and Metabolic Phenotypes of Polycystic Ovary Syndrome in Female Mice. Biol Reprod 93 69.

Kim NR, David K, Corbeels K, Khalil R, Antonio L, Schollaert D, Deboel L, Ohlsson C, Gustafsson JA, Vangoitsenhoven R, et al. 2021 Testosterone Reduces Body Fat in Male Mice by Stimulation of Physical Activity Via Extrahypothalamic ERalpha Signaling. Endocrinology 162.

Krause WC, Rodriguez R, Gegenhuber B, Matharu N, Rodriguez AN, Padilla-Roger AM, Toma K, Herber CB, Correa SM, Duan X, et al. 2021 Oestrogen engages brain MC4R signalling to drive physical activity in female mice. Nature.

Lee YH, Petkova AP, Mottillo EP & Granneman JG 2012 In vivo identification of bipotential adipocyte progenitors recruited by beta3-adrenoceptor activation and high-fat feeding. Cell Metab 15 480–491.

Liu E, Samad F & Mueller BM 2013 Local adipocytes enable estrogen-dependent breast cancer growth: Role of leptin and aromatase. Adipocyte 2 165–169.

Lubahn DB, Moyer JS, Golding TS, Couse JF, Korach KS & Smithies O 1993 Alteration of reproductive function but not prenatal sexual development after insertional disruption of the mouse estrogen receptor gene. Proc Natl Acad Sci U S A 90 11162–11166.

MacDonald PC, Madden JD, Brenner PF, Wilson JD & Siiteri PK 1979 Origin of estrogen in normal men and in women with testicular feminization. J Clin Endocrinol Metab 49 905–916.

Marin P & Arver S 1998 Androgens and abdominal obesity. Baillieres Clin Endocrinol Metab 12 441–451.

Meyer AS 1955 Conversion of 19-hydroxy-delta 4-androstene-3,17-dione to estrone by endocrine tissue. Biochim Biophys Acta 17 441–442.

Meyer AS, Hayano M, Lindberg MC, Gut M & Rodgers OG 1955 The conversion of delta 4-androstene-3,17-dione-4-C14 and dehydroepiandrosterone by bovine adrenal homogenate preparations. Acta Endocrinol (Copenh) 18 148–168.

Monjo M, Pujol E & Roca P 2005 alpha2-to beta3-Adrenoceptor switch in 3T3-L1 preadipocytes and adipocytes: modulation by testosterone, 17beta-estradiol, and progesterone. Am J Physiol Endocrinol Metab 289 E145–150.

Morishima A, Grumbach MM, Simpson ER, Fisher C & Qin K 1995 Aromatase deficiency in male and female siblings caused by a novel mutation and the physiological role of estrogens. J Clin Endocrinol Metab 80 3689–3698.

Musatov S, Chen W, Pfaff DW, Mobbs CV, Yang XJ, Clegg DJ, Kaplitt MG & Ogawa S 2007 Silencing of estrogen receptor alpha in the ventromedial nucleus of hypothalamus leads to metabolic syndrome. Proc Natl Acad Sci U S A 104 2501–2506.

Muzumdar MD, Tasic B, Miyamichi K, Li L & Luo L 2007 A global double-fluorescent Cre reporter mouse. Genesis 45 593–605.

Nelson LR & Bulun SE 2001 Estrogen production and action. J Am Acad Dermatol 45 S116–124.

Ohlsson C, Hellberg N, Parini P, Vidal O, Bohlooly YM, Rudling M, Lindberg MK, Warner M, Angelin B & Gustafsson JA 2000 Obesity and disturbed lipoprotein profile in estrogen receptor-alpha-deficient male mice. Biochem Biophys Res Commun 278 640–645.

Ohlsson C, Hammarstedt A, Vandenput L, Saarinen N, Ryberg H, Windahl SH, Farman HH, Jansson JO, Moverare-Skrtic S, Smith U, et al. 2017 Increased adipose tissue aromatase activity improves insulin sensitivity and reduces adipose tissue inflammation in male mice. Am J Physiol Endocrinol Metab 313 E450–E462.

Pedersen SB, Kristensen K, Hermann PA, Katzenellenbogen JA & Richelsen B 2004 Estrogen controls lipolysis by up-regulating alpha2A-adrenergic receptors directly in human adipose tissue through the estrogen receptor alpha. Implications for the female fat distribution. J Clin Endocrinol Metab 89 1869–1878.

Polari L, Yatkin E, Martinez Chacon MG, Ahotupa M, Smeds A, Strauss L, Zhang F, Poutanen M, Saarinen N & Makela SI 2015 Weight gain and inflammation regulate aromatase expression in male adipose tissue, as evidenced by reporter gene activity. Mol Cell Endocrinol 412 123–130.

Ramirez I 1980 Relation between estrogen-induced hyperlipemia and food intake and body weight in rats. Physiol Behav 25 511–518.

Rodeheffer MS, Birsoy K & Friedman JM 2008 Identification of white adipocyte progenitor cells in vivo. Cell 135 240–249.

Roesch K, Jadhav AP, Trimarchi JM, Stadler MB, Roska B, Sun BB & Cepko CL 2008 The transcriptome of retinal Muller glial cells. J Comp Neurol 509 225–238.

Schneider G, Kirschner MA, Berkowitz R & Ertel NH 1979 Increased estrogen production in obese men. J Clin Endocrinol Metab 48 633–638.

Sebo ZL & Rodeheffer MS 2021 Testosterone metabolites differentially regulate obesogenesis and fat distribution. Mol Metab 44 101141.

Sebo ZL, Jeffery E, Holtrup B & Rodeheffer MS 2018 A mesodermal fate map for adipose tissue. Development 145.

Seidell JC, Bjorntorp P, Sjostrom L, Kvist H & Sannerstedt R 1990 Visceral fat accumulation in men is positively associated with insulin, glucose, and C-peptide levels, but negatively with testosterone levels. Metabolism 39 897–901.

Shen M & Shi H 2015 Sex Hormones and Their Receptors Regulate Liver Energy Homeostasis. Int J Endocrinol 2015 294278.

Stubbins RE, Holcomb VB, Hong J & Nunez NP 2012 Estrogen modulates abdominal adiposity and protects female mice from obesity and impaired glucose tolerance. Eur J Nutr 51 861–870.

Toth MJ, Tchernof A, Sites CK & Poehlman ET 2000 Effect of menopausal status on body composition and abdominal fat distribution. Int J Obes Relat Metab Disord 24 226–231.

Vermeulen A, Kaufman JM, Goemaere S & van Pottelberg I 2002 Estradiol in elderly men. Aging Male 5 98–102.

Villa A, Della Torre S, Stell A, Cook J, Brown M & Maggi A 2012 Tetradian oscillation of estrogen receptor alpha is necessary to prevent liver lipid deposition. Proc Natl Acad Sci U S A 109 11806–11811.

Wang A, Luo J, Moore W, Alkhalidy H, Wu L, Zhang J, Zhen W, Wang Y, Clegg DJ, Bin X, et al. 2016 GPR30 regulates diet-induced adiposity in female mice and adipogenesis in vitro. Sci Rep 6 34302.

Wang ZV, Deng Y, Wang QA, Sun K & Scherer PE 2010 Identification and characterization of a promoter cassette conferring adipocyte-specific gene expression. Endocrinology 151 2933–2939.

Ward ZJ, Bleich SN, Cradock AL, Barrett JL, Giles CM, Flax C, Long MW & Gortmaker SL 2019 Projected U.S. State-Level Prevalence of Adult Obesity and Severe Obesity. N Engl J Med 381 2440–2450.

Wurtman JJ & Baum MJ 1980 Estrogen reduces total food and carbohydrate intake, but not protein intake, in female rats. Physiol Behav 24 823–827.

Xu Y, Nedungadi TP, Zhu L, Sobhani N, Irani BG, Davis KE, Zhang X, Zou F, Gent LM, Hahner LD, et al. 2011 Distinct hypothalamic neurons mediate estrogenic effects on energy homeostasis and reproduction. Cell Metab 14 453–465.

Yang Y, Smith DL, Jr., Keating KD, Allison DB & Nagy TR 2014 Variations in body weight, food intake and body composition after long-term high-fat diet feeding in C57BL/6J mice. Obesity (Silver Spring) 22 2147–2155.

Zang H, Ryden M, Wahlen K, Dahlman-Wright K, Arner P & Linden Hirschberg A 2007 Effects of testosterone and estrogen treatment on lipolysis signaling pathways in subcutaneous adipose tissue of postmenopausal women. Fertil Steril 88 100–106.

Zhao H, Innes J, Brooks DC, Reierstad S, Yilmaz MB, Lin Z & Bulun SE 2009 A novel promoter controls Cyp19a1 gene expression in mouse adipose tissue. Reprod Biol Endocrinol 7 37.

Zhu L, Brown WC, Cai Q, Krust A, Chambon P, McGuinness OP & Stafford JM 2013 Estrogen treatment after ovariectomy protects against fatty liver and may improve pathway-selective insulin resistance. Diabetes 62 424–434.

