## Supplemental Data 1 for "Insights of the role of estrogen in obesity from two models of ERα deletion"

SUPPLEMENTAL FIGURE 1

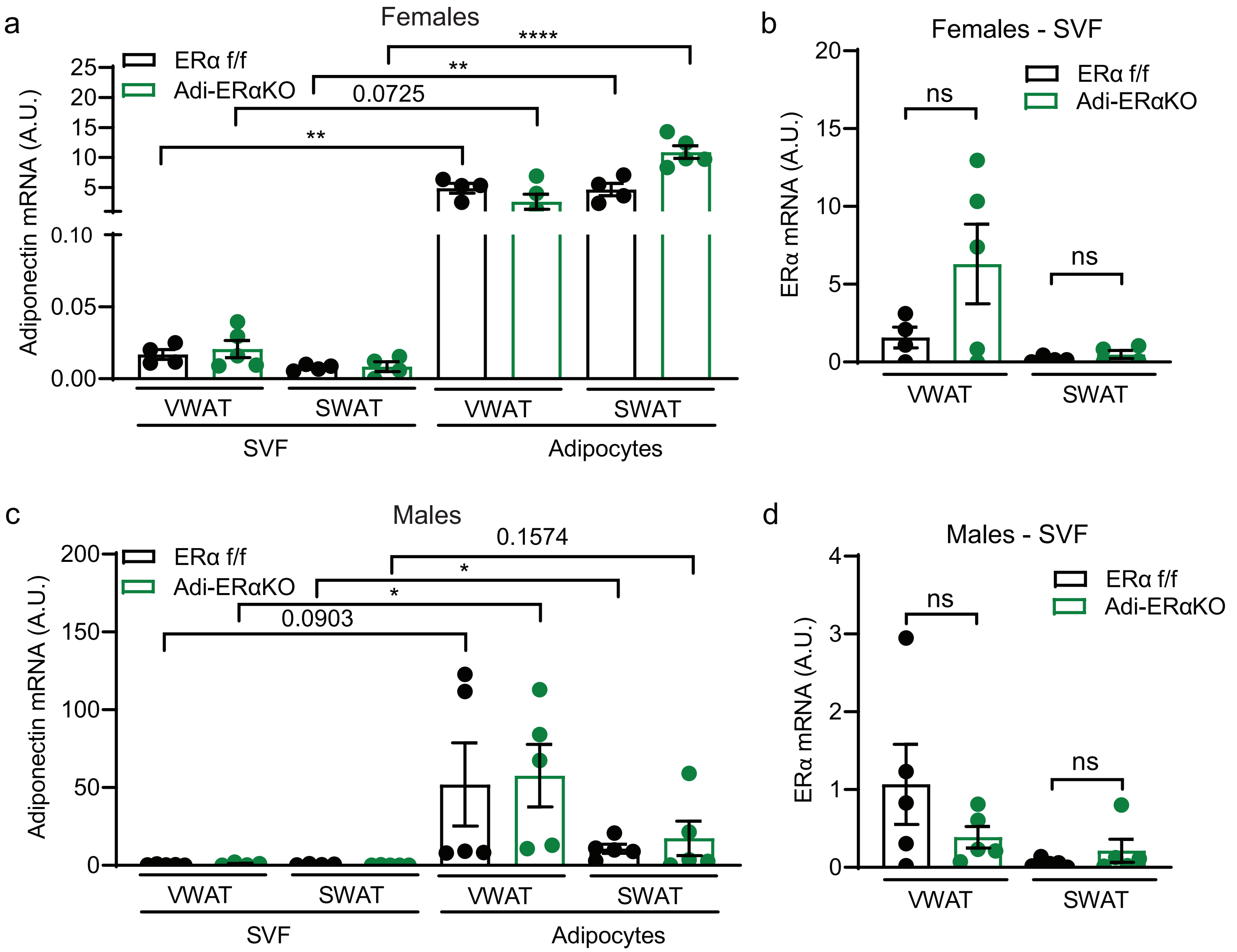

SUPPLEMENTAL FIGURE 2

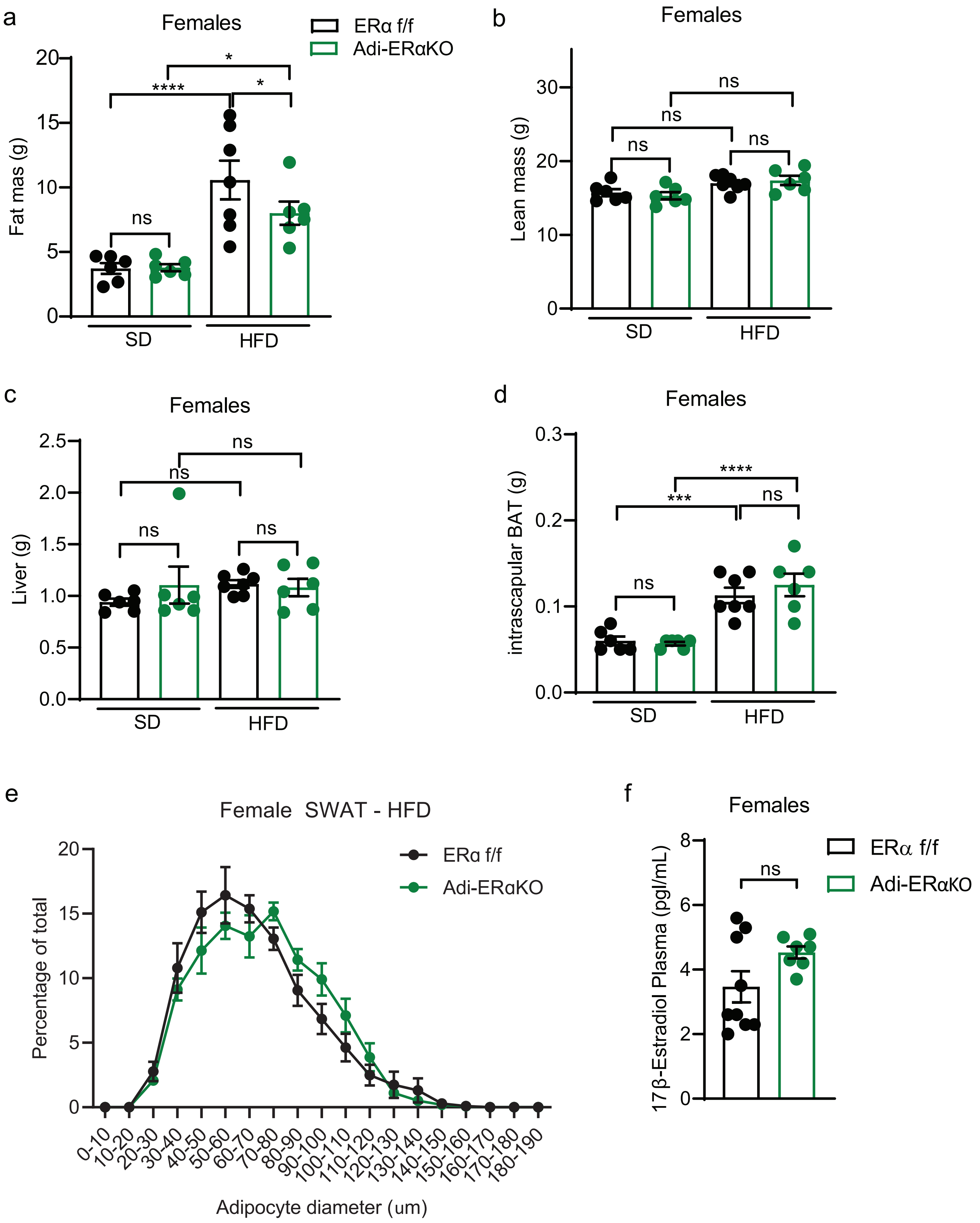

SUPPLEMENTAL FIGURE 3

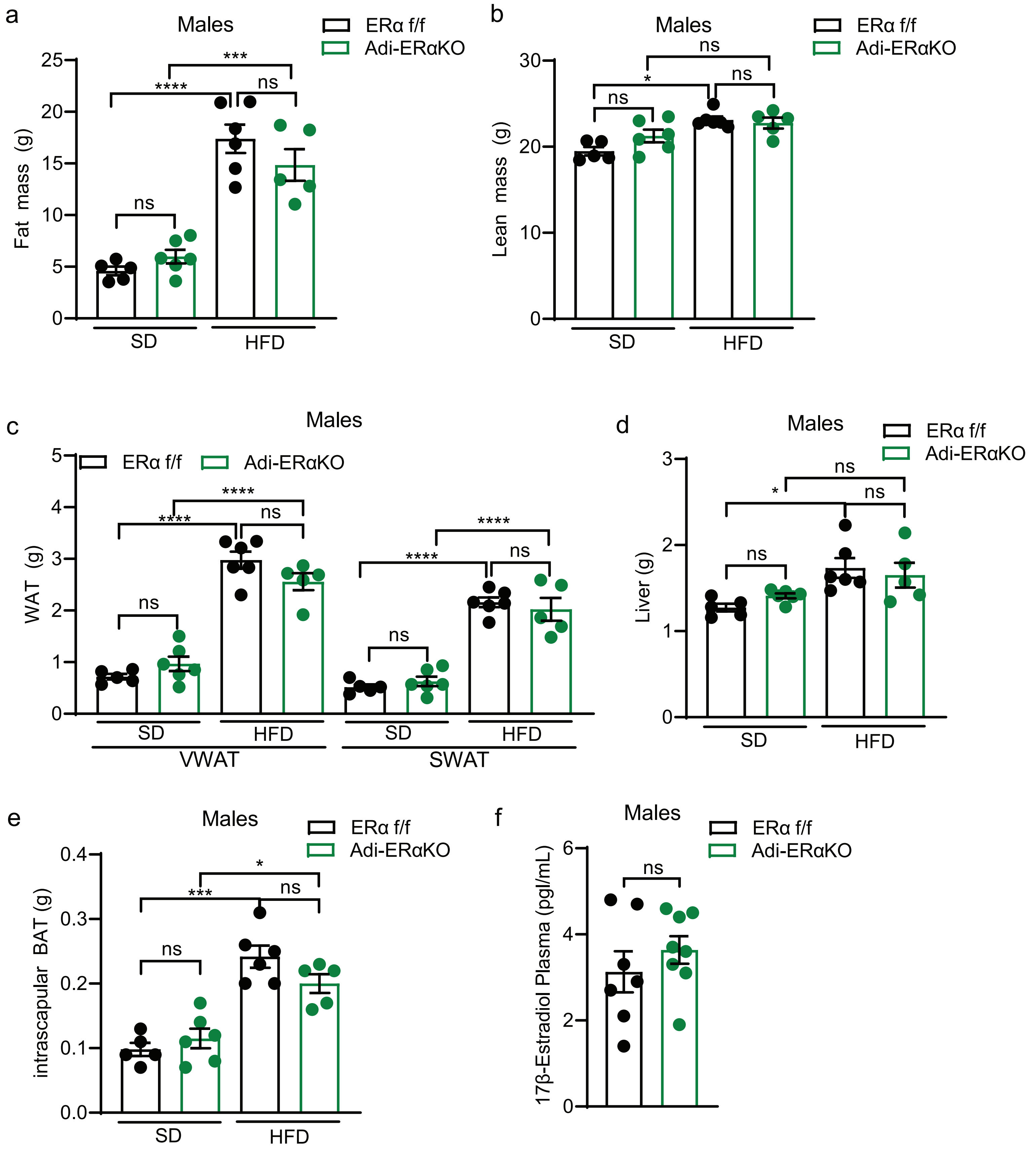

### SUPPLEMENTAL FIGURE 4

a

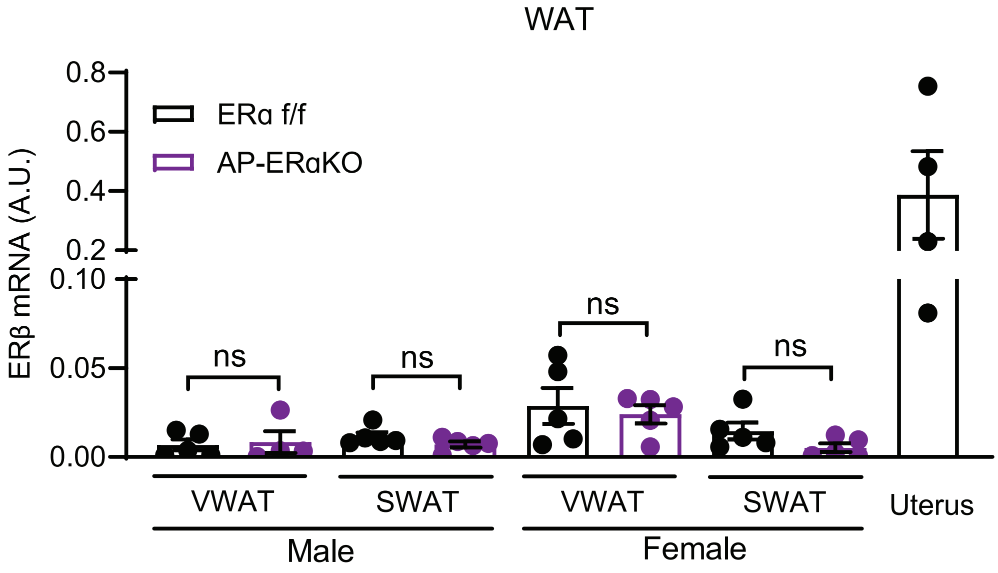

### SUPPLMENTAL FIGURE 5

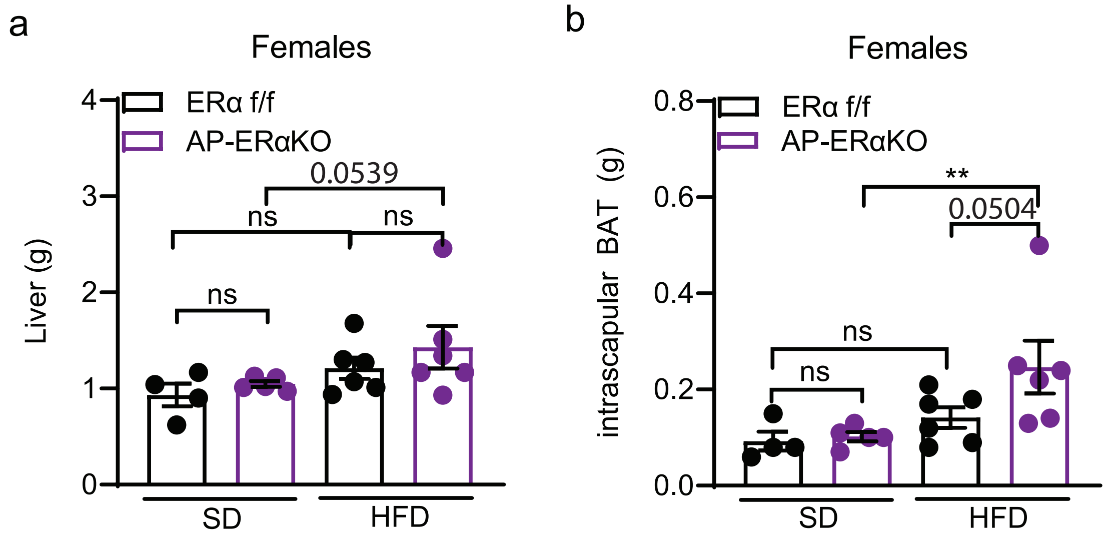

### SUPPLEMENTAL FIGURE 6

a

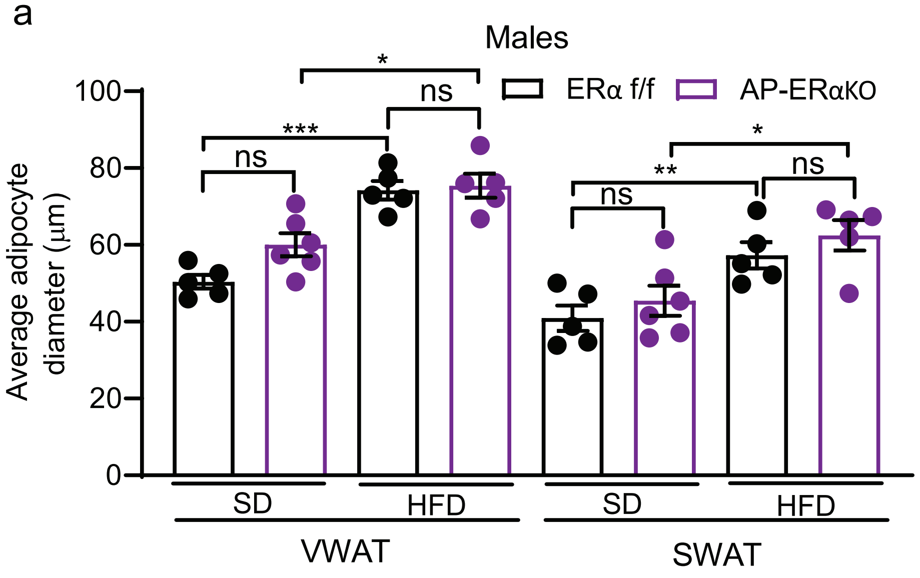


b

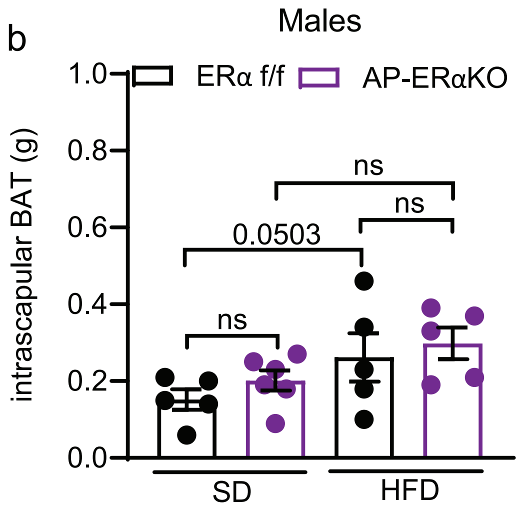


c

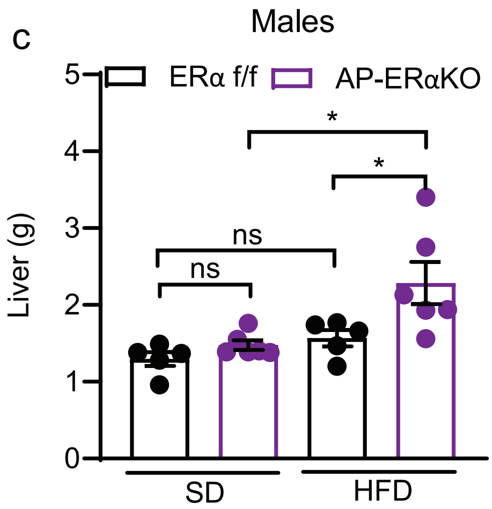

### SUPPLEMENTAL FIGURE 7

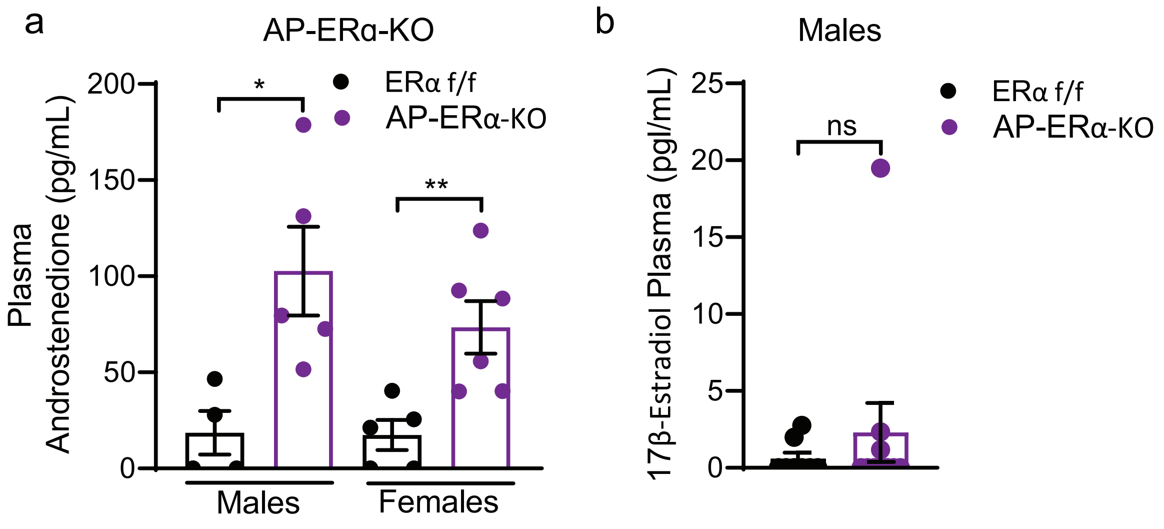

### SUPPLEMENTAL FIGURE 8

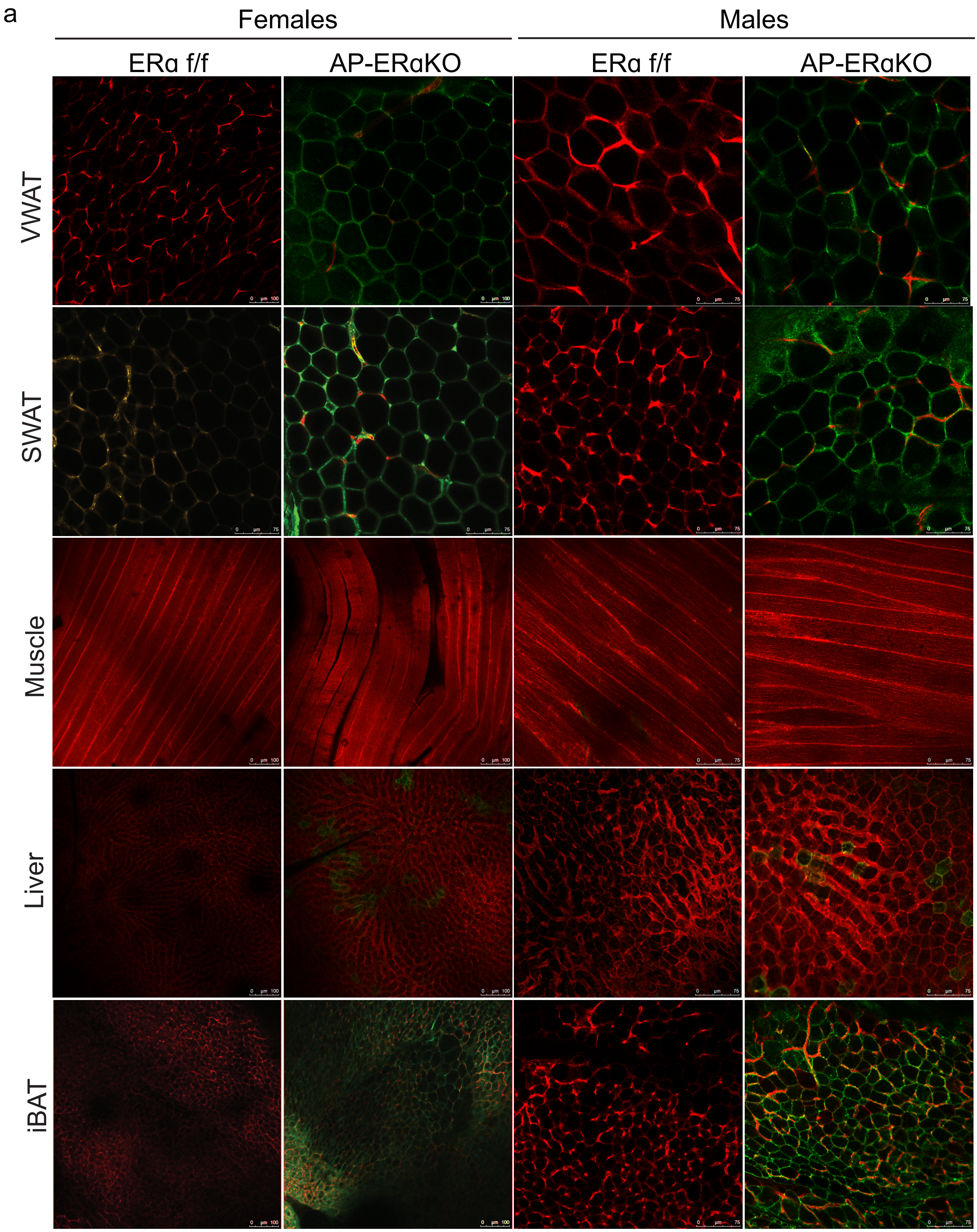

### SUPPLEMENTAL FIGURE 9

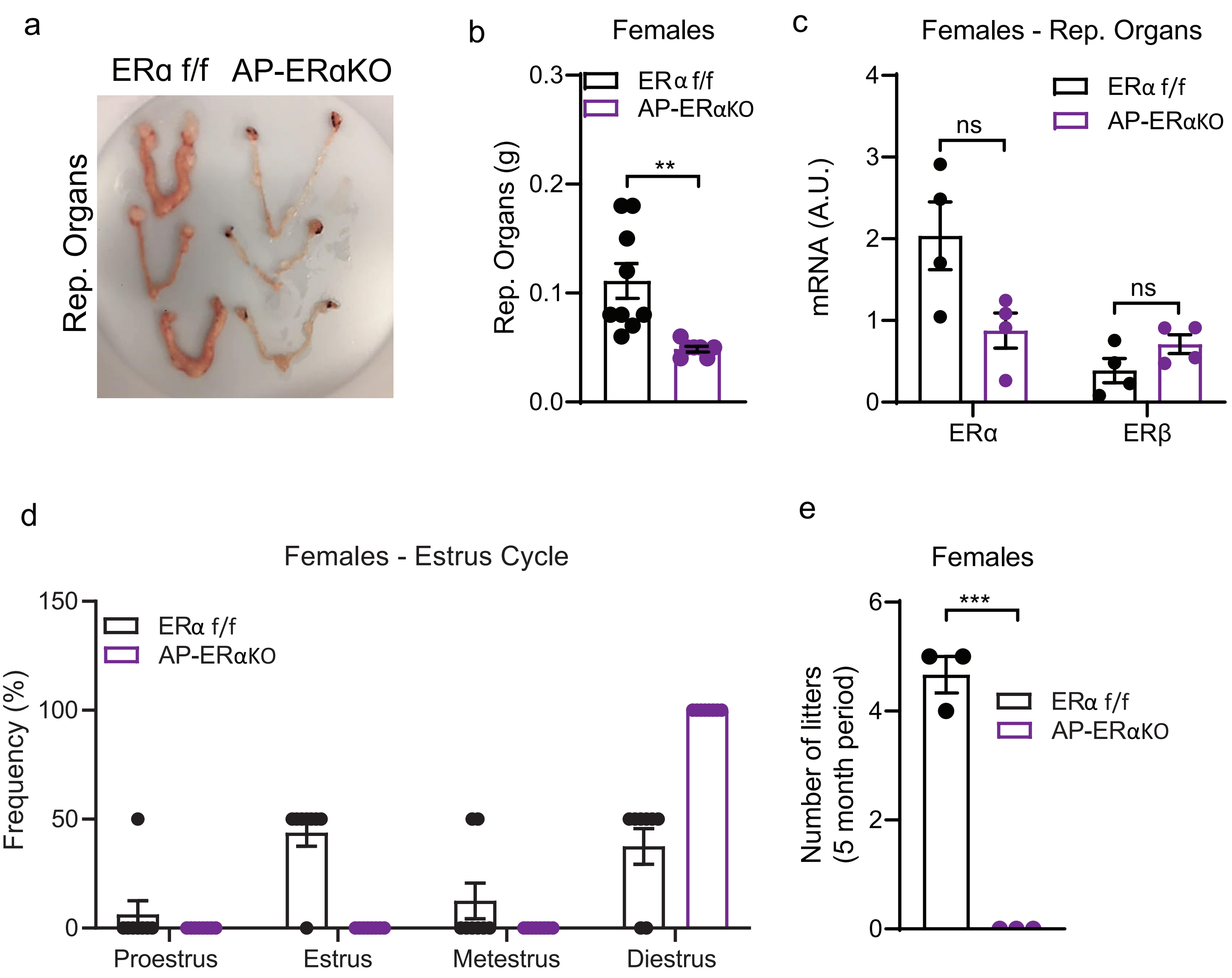
