## Supplemental Data 2 for "Insights of the role of estrogen in obesity from two models of ERα deletion"

**Supplemental Figure Legends**

**Supplemental Figure 1: ERα transcripts in stromal vascular fraction of Adi-ERαKO mice**

(A) Adiponectin expression in stromal vascular fraction (SVF) and mature adipocytes of Adi-ERα-KO females and controls. (n=4 mice per group)
(B) ERα expression in SVF of Adi-ERα-KO females and controls. (n=4-5 mice per group)
(C) Adiponectin expression in SVF and mature adipocytes of Adi-ERα-KO males and controls. (n=5 mice per group)
(D) ERα expression in SVF of Adi-ERα-KO males and controls. (n=5 mice per group)

Statistical significance determined by unpaired t-tests. Error bars represent mean ± S.E.M. * p<0.05, **p<0.01, ****p<0.0001. SVF: stromal vascular fraction, VWAT: visceral fat, SWAT: subcutaneous fat.

**Supplemental Figure 2: Tissue weight of Adi-ERα-KO female mice.**

(A) Total fat mass of female Adi-ERα-KOs and controls at 8 weeks of SD or HFD. (n=6-7 mice per group)
(B) Total lean mass of male Adi-ERα-KOs and controls at 8 weeks of SD or HFD. (n=6-7 mice per group)
(C) Liver weight of female Adi-ERα-KOs and controls at 8 weeks of SD or HFD. (n=6-7 mice per group)
(D) Intrascapular brown adipose tissue (iBAT) weight of female Adi-ERα-KOs and controls at 8 weeks of SD or HFD. (n=6-7 mice per group)
(E) Distribution of adipocyte size (diameter) in SWAT of female Adi-ERα-KOs and controls after 8 weeks of HFD. (n=6-7 mice per group)
(F) Plasma 17-β estradiol levels in Adi-ERα-KO females and controls. (n=7-9 mice per group)

Statistical significance determined by two-way ANOVA with Tukey test for panels A-D. Statistical significance determined by multiple unpaired t-tests for panel E. Statistical significance determined by unpaired t-tests for panel F. Error bars represent mean ± S.E.M. * p<0.05, ***p<0.001, ****p<0.0001. SD: standard diet, HFD: high-fat diet, SWAT: subcutaneous fat.

**Supplemental Figure 3: Lack of ERα in adipocytes does not impact male WAT accumulation.**

(A) Total fat mass of male Adi-ERα-KO males and controls after 8 weeks of SD and HFD. (n=5-6 mice per group)
(B) Total lean mass of male Adi- ERα-KO males and controls after 8 weeks of SD and HFD. (n=5-6 mice per group)
(C) VWAT and SWAT accumulation of male Adi-ERα-KO and controls after 8 weeks of SD and HFD. (n=5-6 mice per group)
(D) Liver weight of male Adi-ERα-KOs and controls at 8 weeks of SD or HFD. (n=5-6 mice per group)
(E) Intrascapular brown adipose tissue (iBAT) weight of male Adi-ERα-KOs and controls at 8 weeks of SD or HFD. (n=5-6 mice per group)
(F) Plasma 17-β estradiol levels in Adi-ERα-KO males and controls. (n=7-8 mice per group)

Statistical significance determined by two-way ANOVA with Tukey test for panels A-E and unpaired t-test for panel F. Error bars represent mean ± S.E.M. *p<0.05, ***p<0.001, ****p<0.0001. SD: standard diet, HFD: high-fat diet, VWAT: visceral fat, SWAT: subcutaneous fat.

**Supplemental Figure 4: Expression of ERβ in AP-ERα-KO mice.**

(A) ERβ expression in WAT from male and female AP-ERα-KO mice. Female uterus included as a positive control. (n=4-5 mice per group)

Statistical significance determined by unpaired t-tests. ns= not significant. Error bars represent mean ± S.E.M. VWAT: visceral fat, SWAT: subcutaneous fat.

**Supplemental Figure 5: Tissues weights of female AP-ERα-KO.**

(A) Liver weight of female AP-ERα-KOs and controls at 8 weeks of SD or HFD. (n=5-6 mice per group)
(B) Intrascapular brown adipose tissue (iBAT) weight of female AP-ERα-KOs and controls at 8 weeks of SD or HFD. (n=5-6 mice per group)

Statistical significance determined by two-way ANOVA with Tukey test. Error bars represent mean ± S.E.M. **p<0.01. SD: standard diet, HFD: high-fat diet.

**Supplemental Figure 6: Male AP-ERα-KO mice don’t have bigger adipocytes on a HFD.**

(A) Average adipocyte diameter (µm) of male AP-ERα-KO and controls after 8 weeks of SD and HFD. (n=5-6 per group)
(B) Intrascapular brown adipose tissue (iBAT) weight of male AP-ERα-KOs and controls after 8 weeks of SD or HFD. (n=5-6 mice per group)
(C) Liver weight of male AP-ERα-KOs and controls after 8 weeks of SD or HFD. (n=5-6 mice per group)

Statistical significance determined by two-way ANOVA with Tukey test. Error bars represent mean ± S.E.M. *p<0.05, **p<0.01, ***p<0.001, ****p<0.0001. SD: standard diet, HFD: high-fat diet, VWAT: visceral fat, SWAT: subcutaneous fat.

**Supplemental Figure 7: Altered sex hormones in plasma of AP-ERα-KO mice.**

(A) Plasma androstenedione of male and female AP-ERα-KO mice. (n=4-6 mice per group)
(B) Plasma 17-β estradiol of male AP-ERα-KO mice. (n=8-10 mice per group)

Statistical significance determined by unpaired t-tests. Error bars represent mean ± S.E.M. * p<0.05, **p<0.01.

**Supplemental Figure 8: PdgfRα labels WAT and iBAT in male and female mice.**

(A) Whole-mount confocal images of AP-ERα-KO; mTmG tissues taken at 20X. GFP fluorescence indicates presence of cre activity. Scale bar is 100µm. (n=1 mice per group)

VWAT: visceral fat, SWAT: subcutaneous fat, iBAT: intrascapular brown adipose tissue.

**Supplemental Figure 9: Female AP-ERα-KO mice are infertile.**

(A) Gross morphology of female reproductive organs from AP-ERα-KO females and controls. (n=3 mice per group)
(B) Weight of reproductive organs from female AP-ERα-KO and controls. (n=7-9 mice per group)
(C) Expression of ERα and ERβ from reproductive organs of female AP-ERα-KO and controls. (n=4 mice per group)
(D) Frequency of each estrus cycle stage in female AP-ERα-KO and controls. (n= 6-7 mice per group)
(E) Litter numbers of female AP-ERα-KO or controls when breeding with wildtype males for 5 months. (n=3 female mice per group)

Statistical significance determined by unpaired t-tests. Error bars represent mean ± S.E.M. **p<0.01, ***p<0.001.
